# Meta-omics reveals role of photosynthesis in Microbially Induced Carbonate Precipitation at a CO_2_-rich Geyser

**DOI:** 10.1101/2024.07.20.604422

**Authors:** Marlene J. Violette, Ethan Hyland, Landon Burgener, Adit Ghosh, Brina M. Montoya, Manuel Kleiner

## Abstract

Microbially induced carbonate precipitation (MICP) is a natural process with potential biotechnological applications to address both carbon sequestration and sustainable construction needs. However, our understanding of the microbial processes involved in MICP is limited to a few well-researched pathways such as ureolytic hydrolysis. To expand our knowledge of MICP, we conducted an omics-based study on sedimentary communities from travertine around the CO_2_-driven Crystal Geyser near Green River, Utah. Using metagenomics and metaproteomics, we identified the community members and potential metabolic pathways involved in MICP. We found variations in microbial community composition between the two sites we sampled, but *Rhodobacterales* were consistently the most abundant order, including both chemoheterotrophs and anoxygenic phototrophs. We also identified several highly abundant genera of *Cyanobacteriales*. The dominance of these community members across both sites and the abundant presence of photosynthesis-related proteins suggest that photosynthesis could play a role in MICP at Crystal Geyser. We also found abundant bacterial proteins involved in phosphorous starvation response at both sites suggesting that P-limitation shapes both composition and function of the microbial community driving MICP. Our study emphasizes the possible involvement of photosynthesis in MICP processes at Crystal Geyser and suggests that P-limitation may either hinder or facilitate MICP within the community.

## Introduction

Carbonate deposits are ubiquitous on Earth, accounting for one-sixth of global sedimentary rocks (1). A major part of these carbonate deposits is of microbial origin (2). Microbially induced carbonate precipitation (MICP) refers to the biogeochemical process in which calcium carbonate is precipitated by altering the local geochemical environment (3). These alterations occur as a by-product of common microbial metabolic activities that lead to increased local carbonate content and pH, thereby saturating the solution with respect to carbonate. If supersaturation is reached and Ca^2+^ is present, CaCO_3_ will precipitate out of solution. Several microbial metabolic pathways have been proposed to be involved in MICP including photosynthesis (4), methane oxidation (5), ureolysis (3), denitrification (6), ammonification (7) and sulfate reduction (8). In addition, cell surface structures such as extracellular polymeric substances (EPS) (9) and S-layer proteins (10) have been linked to MICP due to their capability of binding positively charged ions such as Ca^2+^ and thus serving as crystal nucleus for carbonate precipitation, thereby entrapping the microbe.

Many studies in the realm of biomineralization and MICP have focused on organo-sedimentary structures such as stromatolites. Stromatolites are of particular interest for astro-microbiologists as fossilized stromatolites are the oldest macrofossil records of life on Earth. While stromatolite morphology and taxonomic composition of the microbial communities vary, they always feature steep physico-chemical gradients and almost complete microscale cycles of C, N, and S (11). These gradients are very similar to the nutrient gradients commonly found in soils and sediments and thus might serve as analogs for biomineralization communities in these environments. In recent years, MICP has become the focus of cross-disciplinary studies in the fields of geotechnical and structural engineering as it holds the potential to become a major player in bio-mediated technologies such as carbon sequestration and geological carbon storage (12), bioremediation (13), concrete restoration, or soil reinforcement (14). While a tremendous amount of work has been done on MICP in the context of these applications, several challenges remain. For instance, most application-oriented studies focus on ureolytic bacteria using *Sporosarcina pasteurii* as a model organism due its high urease activity, presence of EPS, and weak motility (15). However, urea hydrolysis results in ammonia as a product, which is unfavorable for many applications due its corrosiveness and toxicity (16). The applicability of other mechanisms for MICP is heavily understudied and some metabolic mechanisms are still debated (e.g., sulfate reduction) and need to be evaluated on a case-to-case basis. Moreover, most studies in this field are conducted in axenic culture, which does not reflect environmental conditions and community dynamics properly and may thus not correspond to conditions encountered during field deployment.

To better understand the microbial ecology of MICP in natural environments, we chose to evaluate microbial communities derived from travertine adjacent to Crystal Geyser (CG), which is located near Green River, Utah. CG is a cold-driven, CO_2_ rich geyser, which is surrounded by colorful travertine that has been suggested to be generated through MICP (17–19). We chose CG sediments because previous studies found CaCO_3_ polymorphism throughout the travertine and entrapped microbial structures within the mineral crystals supporting the idea of a microbial influence in travertine formation. We used a cultivation-independent, multi-omics approach combined with geochemical measurements to investigate metabolic pathways and physiologies potentially involved in MICP at CG.

## Material and Methods

### Sampling

We collected samples from the top 20 cm of travertine adjacent to Crystal Geyser, Utah in November 2019 and June 2021 (38.9384° N, 110.1354° W) wearing gloves at all times. We recovered travertine samples using hammers and tweezers that were autoclaved and baked out at 450°C for 1 hour before sampling. We sampled 1 m away from the borehole (CG-1) and 10 m away from the borehole (CG-10). Gloves were changed and sampling equipment was cleaned with 70% ethanol before switching to the CG-10 site. We preserved all collected samples in RNAlater-like solution (20) in a 1:10 sediment: RNAlater-like solution ratio which has been verified for metagenomic and metaproteomics analysis in previous work (21,22). We transported all samples at ambient temperature to North Carolina State University (Raleigh, USA). In the laboratory, we removed the preservation solution by centrifugation at 14,000 × g for 10 min and ground and homogenized the travertine with mortars and pestles from which organic carbon had been removed by baking at 450°C for 1 hour. Homogenates were aliquoted into 50 mL falcon tubes and immediately frozen at −80°C until further analysis.

### Carbonate quantification

To quantify the CaCO_3_ content we used a gasometric method described by O’Toole et al. (23). We generated a calibration curve using pure 100% CaCO_3_ (VWR International) and 1 M Hydrochloric acid (HCl) (VWR International). We determined the CaCO_3_ content of Crystal Geyser sediments using roughly 13 g of ground travertine homogenate per replicate. As a reference, we also quantified CaCO_3_ of stromatolite samples that were collected in 2019 at the Rife Bed outside of Eden, Wyoming (41.9663° N, 109.2501° W) and sandy soils collected in 2021 near Green River, Utah (38.801335° N, −109.945159° E).

### Isotope-ratio mass spectrometry

We carried out isotope ratio mass spectrometry on organic carbon and carbonates as described in the Supplementary Methods.

### Scanning electron microscopy and energy-dispersive X-ray spectrometry

We carried out scanning electron microscopy and energy-dispersive X-ray spectrometry as described in the Supplementary Methods.

### DNA extraction and sequencing

We lysed 250 mg of each travertine homogenate (2 replicates per site) by bead-beating in lysing matrix Y tubes (MP Biomedicals) with a Bead Ruptor Elite (Omni International) using 6 cycles of 30 s at 3.25 m/s with 1-min dwell time between cycles. DNA was extracted in two technical replicates per site using the DNeasy PowerSoil Pro Kit (Qiagen) according to the manufacturer’s instructions. We determined DNA concentrations using the Qubit™ dsDNA HS assay (Thermo Fisher Scientific) and combined DNA extracts of each technical replicate per site. We aliquoted DNA extracts and sent equal amounts of extracts for shotgun metagenomic sequencing at two different sequencing centers. Metagenomic library preparation and DNA sequencing were conducted at the Genomic Science Laboratory (GSL) at North Carolina State University and the Beijing Genomics Institute (BGI). At GSL, the TruSeq® Nano DNA Library Prep Kit (Illumina) was used for library preparation and DNA sequencing was performed using the Illumina NovaSeq6000 sequencer (150 paired end, SP 150 PE lane (117,153,983 reads for CG-1 and 113,371,508 reads for CG-10)). At BGI, the MGIEasy Universal DNA Prep Set (MGI) was used for library preparation and DNA sequencing was performed using the DNBSEQ sequencer (150 paired end, SP 150 PE lane (111,779,984 reads for CG-1 and 112,180,021 reads for CG-10)).

### Assembly, binning and annotation of metagenomes

We assessed raw sequence quality using FastQC (24). We decontaminated raw reads targeting phi X174 using bbsplit and trimmed adapters with bbduk from the bbmap suite (25). Read error correction was performed using BayesHammer in MetaSpades (26). We conducted a taxonomic raw read assessment using phyloflash (27). We then co-assembled trimmed and error corrected reads using libraries from both sequencing facilities (GSL and BGI) of each sample respectively. Assemblies were performed using MEGAHIT (28) with the following parameters: --k-min 33 --k-max 127 --k-step 10 --mem-flag 0. To separate eukaryotic and prokaryotic contigs, we classified contigs with Whokaryote (29) using default parameters. All contigs that were identified as either prokaryotic or eukaryotic were then handled separately as described in the supplemental methods. We manually recovered all contigs that were below whokayrote’s 5000 bp threshold and were thus filtered out. We will refer to these contigs as “unassigned contigs” throughout this paper. Genes on the unassigned contigs were predicted using prodigal (30) with default settings and used for protein database construction (Fig. S1). Prokaryotic and eukaryotic sequences were handled as described in the Supplementary methods.

### Bacterial community composition and phylogenomics

Bacterial and archaeal genomes were taxonomically classified using GTDB-Tk (31) with the classify workflow using default settings. We selected all bins that were at least of medium quality as defined by Bowers et al. (32) for tree construction. We constructed a phylogenomic tree with GToTree v1.6.34 (33), using the integrated universal single copy gene set (SCGs) derived from Hug et al. (34) (16 target genes). Only bins that contained at least 40% of the defined SCGs were used for tree construction. Briefly, prodigal v2.6.3 (30) was used to predict genes on input genomes provided as fasta files. Target genes were identified with HMMER3 v3.2.2 (35), individually aligned with muscle v5.1 (36), trimmed with trimal v1.4.rev15 (37), and concatenated prior to phylogenetic estimation with FastTree2 v2.1.11 (38). We visualized the tree using the interactive tree of life (ITOL) (39).

### Protein extraction, peptide preparation, and determination

We extracted proteins in triplicate per sampling site. We prepared tryptic peptides by combining a trichloroacetic acid (TCA) precipitation protocol adapted from Qian and Hettich (40) and the filter-aided-sample-preparation (FASP) protocol (41) (Supplementary Methods).

### One-dimensional liquid chromatography–tandem mass spectrometry

All samples were analyzed by one-dimensional liquid chromatography-tandem mass spectrometry (1D-LC-MS/MS). For each sample, 900 ng of tryptic peptides were loaded with an UltiMate 3000 RSLCnano liquid chromatograph (Thermo Fisher Scientific) in loading solvent A (2% acetonitrile, 0.05% trifluoroacetic acid) onto a 5-mm, 300-μm-inner diameter C18 Acclaim PepMap100 pre-column and desalted (Thermo Fisher Scientific). Peptides were then separated on a 75-cm × 75-μm analytical EASY-Spray column packed with PepMap RSLC C18, 2-μm material (ES905, Thermo Fisher Scientific) heated to 60°C via the integrated column heater at a flow rate of 300 nL min−1 using a 140-min gradient going from 95% buffer A (0.1% formic acid) to 31% buffer B (0.1% formic acid, 80% acetonitrile) in 102 min, then to 50% B in 18 min, to 99% B in 1 min, and ending with 99% B. Carryover was reduced by wash runs (injection of 20 μl acetonitrile with 99% eluent buffer B) between samples. The analytical column was connected to an Orbitrap Exploris™ 480 Mass Spectrometer (Thermo Fisher Scientific) via an Easy-Spray source. Eluting peptides were ionized via electrospray ionization (ESI). MS^1^ spectra were acquired by performing a full MS scan at a resolution of 60,000 on a 380 to 1,600 m/z window. MS^2^ spectra were acquired using a data-dependent approach by selecting for fragmentation the 15 most abundant ions from the precursor MS^1^ spectra. A normalized collision energy of 27% was applied in the higher-energy collisional dissociation (HCD) cell to generate the peptide fragments for MS^2^ spectra. Other settings of the data-dependent acquisition included a maximum injection time of 50 ms, a dynamic exclusion of 25 s, and exclusion of ions of +1 charge state from fragmentation. On average, 82,000 MS/MS spectra were acquired per sample.

### Proteinaceous biomass calculations

We used a method adapted from Kleiner et al. (42) to calculate proteinaceous biomass with the following modification. Instead of using FidoCT for protein inference in Proteome Discoverer and filtering for proteins with at least 2 protein unique peptides (PUPs), we used the Protein Validator node for protein inference and filtered for proteins with at least 2 PUPs.

### Data availability

The metagenomic raw reads have been deposited to Bioproject (PRJNA961082).The metaproteomics MS data and protein sequence database have been deposited to the ProteomeXchange Consortium via the PRIDE partner repository (43), with the following data set identifiers: PXD041379.

## Results and discussion

We studied microbial communities in Crystal Geyser (CG) travertine to understand their role in travertine formation. Using a metagenomics-metaproteomics approach and stable isotope analysis, we aimed to characterize microbial metabolisms and physiology. Samples were taken 1 m (CG-1) and 10 m (CG-10) from the borehole to see how proximity to the CO_2_-enriched water source affects microbial communities and activities related to carbonate precipitation.

### Minor differences in CaCO_3_ quantities and elemental composition in travertine collected from Crystal Geyser

Our scanning electron microscopy (SEM) and energy-dispersive X-ray spectrometry (EDX) analyses showed the presence of two CaCO_3_ polymorphs at CG-1, which appear to be aragonite blades and cuboidal calcite crystal (Fig 1D, Figure S2-4). This is consistent with previous reports that Crystal Geyser waters are supersaturated with respect to both aragonite and calcite (44). We did not find blade-shaped crystals in travertine collected at CG-10 and only observed cuboidal crystals that are likely calcite (Fig. 1E, Figure S5-6).

**Figure 1:**
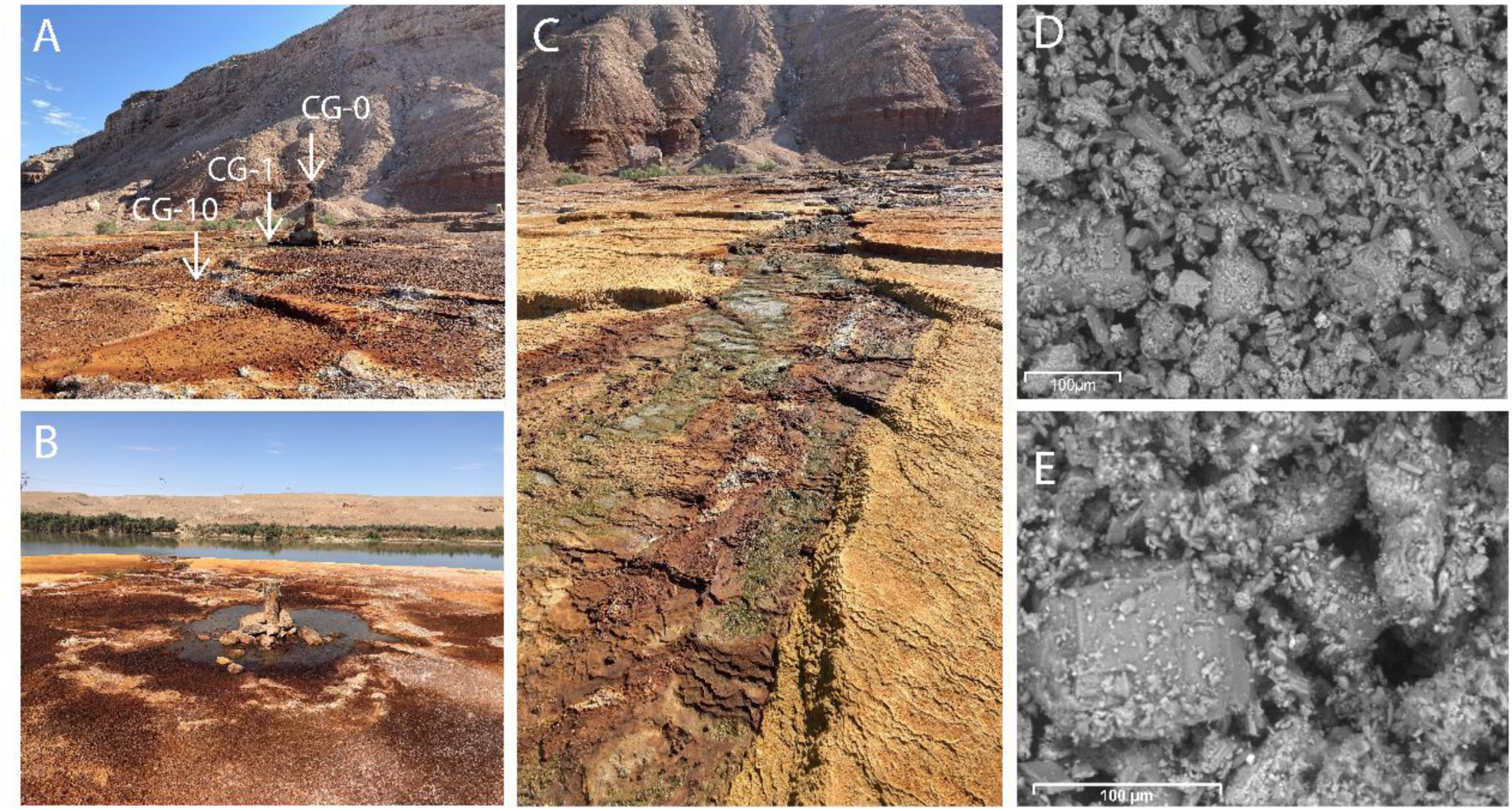
Images of the Crystal Geyser (CG) field site and CaCO_3_ crystals. A) Overview of the CG sampling site. Arrow CG-0 marks the geyser borehole, arrow CG-1 marks sampling site 1 m from borehole and arrow CG-10 marks sampling site 10 m from borehole. B) CG borehole surrounded by a rimpool of geyser water. Green River in the background. C) Flow path of the geyser waters to Green River. D) Scanning electron microscopy (SEM) image of travertine collected at CG-1. E) SEM image of travertine collected at CG-10.

To assess what elements are present in CG travertine and are thus available to the microbial communities, we performed elemental analysis of CG travertine. We found that both sites were similar in elemental composition with oxygen, calcium and carbon being the most abundant (in order of decreasing abundance) (Tables S1 and S2). We also found that both sites contained varying amounts of nitrogen and sulfur; however, these were considerably lower in abundance than oxygen, calcium and carbon. Interestingly, we identified differences in the presence of silicon, magnesium, and sodium between the CG-1 and CG-10 travertine samples. Specifically, silicon was detected in multiple spectra of CG-1 while it was not detected in CG-10 samples. Conversely, small amounts of magnesium and sodium were found in spectra from CG-10 but not in CG-1. These variations could be attributed to a solubility gradient of the geyser discharge, which is known to contain various dissolved ions including Si, Mg, and Na (45). Silica tends to precipitate out first as water evaporates, followed by salts such as Mg and Na, which could explain the differences observed in the two samples (46). Interestingly, our analysis did not detect any phosphorus in any of our samples, including the geyser water that was analyzed in previous work (61). Phosphorus is a crucial element for life, and its absence in the samples suggests that microbial communities at CG might be phosphorus limited. Additionally, we observed significant variations in the total amount of organic carbon between the sites, with CG-1 containing nearly twice as much organic carbon as CG-10 (Table S3). These differences in organic carbon between the two sites might indicate varying sources of organic matter and/or differences in microbial biomass production. In summary, our findings suggest that CG travertine is primarily composed of CaCO_3_, as evidenced by the high abundance of oxygen, calcium, and carbon, and the observed differences in elemental composition between sites may impact microbial community function and biomass production.

We conducted a gasometric analysis to specifically quantify the CaCO_3_ content of the two sampling sites. For comparison with samples rich in CaCO_3_ and those with low amounts of CaCO_3_, we measured the quantity of CaCO_3_ in stromatolites—known for being predominantly composed of CaCO_3_—and sandy desert soils, which typically contain lower amounts of CaCO_3_. We found minor differences in terms of CaCO_3_ quantities between CG-1 and CG-10 (Figure S7, Table S4) and these differences were not significant (Student’s t test, P > 0.05) (Table S5). CaCO_3_ quantities of CG samples (8.38-12.52%) were significantly different from both the high CaCO_3_ sample (13.36-15.73%) and the low CaCO_3_ sample (0.59-1.41%). However, CG samples were on average closer to the CaCO_3_ quantities of the stromatolite samples compared to the soil samples. Our findings confirmed the previously assumed relatively large amounts of CaCO_3_ at CG. Moreover, our analysis revealed that there is no significant difference in CaCO_3_ quantities between the two sampling sites suggesting that the proximity to the geyser borehole may not have a significant impact on the levels of CaCO_3_ precipitation.

### Variations in microbial community composition suggests ecological differences between sites

Metagenomic sequencing of microbial communities recovered from CG travertine revealed differences in taxonomic diversity between the two sampling sites. We recovered 197 bins from CG-1 samples, out of which 132 were at least of medium quality or higher (≥50% complete and ≤10% contamination). In contrast, we obtained 343 bins from CG-10 with 207 medium/high quality bins) (Supplementary File 1). To assess microbial diversity and potential differences in community composition between both sampling sites, we calculated a phylogenomic tree based on 16 ribosomal proteins extracted from all bins that were at least of medium quality using GToTree. Only bins with >40% alignment coverage were retained for the tree analysis resulting in 232 bins remaining for our phylogenetic tree analysis. We identified 78 different microbial orders for both CG-1 and CG-10 combined (Fig. 2). Out of these 78 orders, 76 were bacterial and 2 archaeal, the latter of which were only present at CG-10 (Supplementary File 1). We also identified eukaryotic sequences but they were limited in number and we could not taxonomically identify them beyond the kingdom level.

**Fig. 2:**
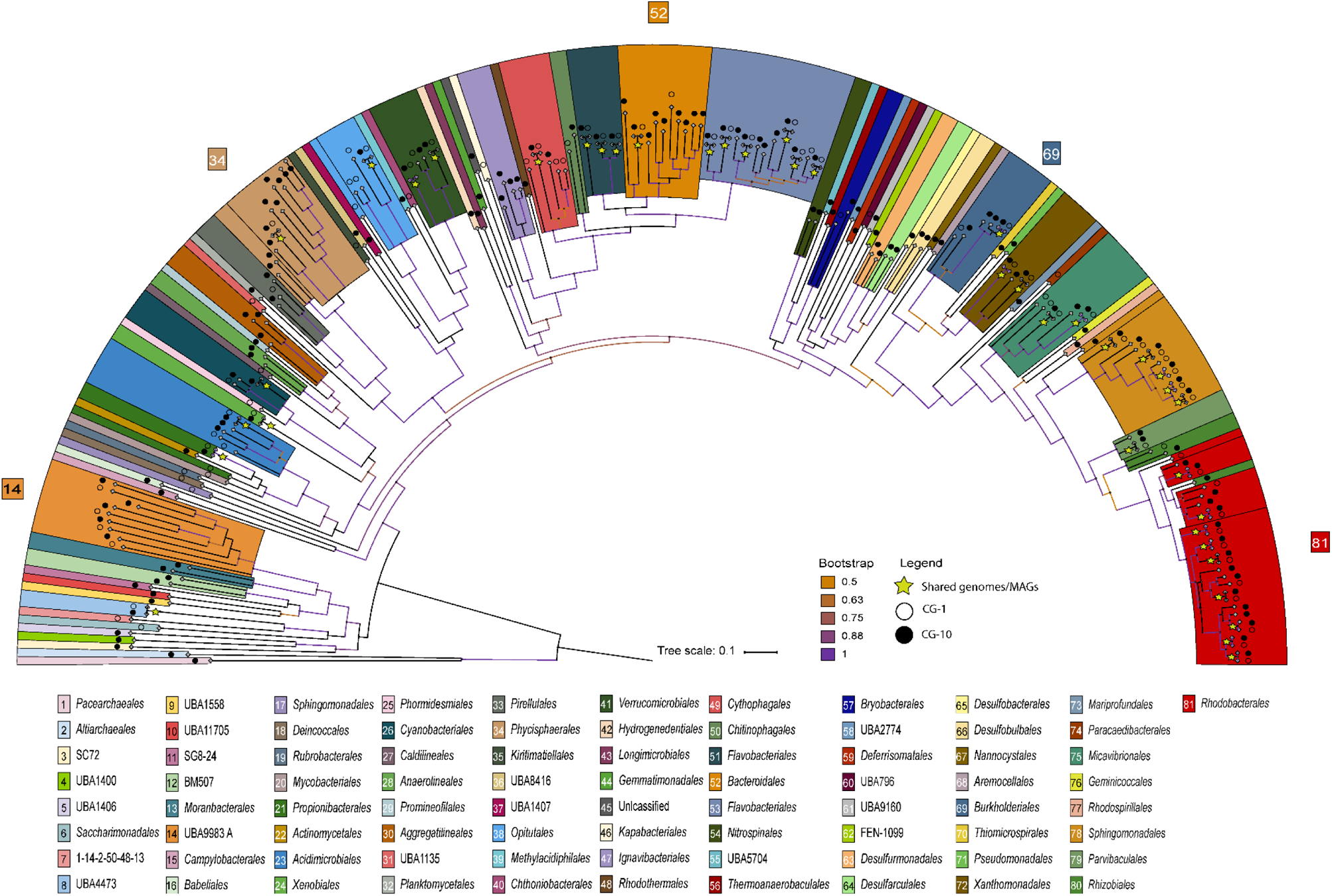
Diversity of bins recovered from CG travertine. The phylogenomic tree was generated based on 16 concatenated ribosomal proteins from 339 medium to high-quality bins using GToTree. Bins were reconstructed for 81 different order-level lineages (107 bins had <40% alignment coverage and are thus not displayed). Taxonomic classifications are according to GTDB-Tk. The scale corresponds to the average amino acid substitution over the alignment.

We found that *Rhodobacterales* was the most diverse order in terms of number of bins that could be classified by GTDB-Tk at the order level. At CG-1, 12 bins out of 84 CG-1 bins that remained in our tree analysis were classified as *Rhodobacterales,* while for CG-10 it was 16 out of 148. We also found large numbers of bins assigned to *Flavobacteriales* (10 at CG-1 and 13 at CG-10), and *Sphingomonadales* (8 at CG-1 and 7 at CG-10) at both sites. Interestingly, we recovered 9 bins that were classified as *Bacteroidales* at CG-10, while we only found 2 bins of this order at CG-1. There were several bacterial orders unique to one sampling site suggesting potential ecological differences between the two sites. For example, we found 12 microbial orders to be unique to CG-1; however, they were not abundant in terms of number of bins as all were represented only by 1 bin (Supplementary File 1). At CG-10, we found 44 orders that were unique to CG-10. Out of these orders, the most abundant in terms of numbers of recovered bins were *Ignavibacteriales* and *Aggregatilineales*, with 4 and 3 bins respectively. *Ignavibacteriales*, specifically from the *Melioribacteracea*e family, were first found in a hot water microbial mat from a Russian oil well and have been described to be facultative anaerobe, obligate chemoorganotrophs (47). Initially thought to belong to *Chlorobi*, they were proposed to form a separate phylum, *Ignavibacteriae*, though this is debated (47,48). Probst et al. (61) also found *Melioribacteraceae* in CG groundwater, thus we would have expected them at CG-1 due to borehole proximity. *Aggregatilineales*, recently proposed as their own order in the phylum of phototrophic *Chloroflexota*, were found in extreme environments and linked to biofilm formation (49,50). Since *Ignavibacteriales* and *Aggregatilineales* were only recovered at CG-10, we speculate they occupy a CG-10-specific niche possibly contributing to travertine formation. Moreover, we found 2 bins of each *Anaerolineales, Desulfobulbales*, *Desulfarculales*, *Desulfuromonadales,* taxonomically unassigned *Patescibactria* (order ID BM507 according to GTDB-Tk), and *Moranbacterales* (*Patescibacteria*) unique to CG-10. Interestingly, we found 8 different orders of *Patescibacteria* unique to CG-10, most of which were only represented by 1 bin and taxonomically unassigned beyond the class level. This was particularly surprising as previous work reported several MAGs of the superphylum *Patescibacteria*, previously known as *Candidate Phyla Radiation* (CPR) bacteria, to be present at deep and intermediate depths in the water of the geyser itself (45,51,52). Based on this, we would have expected to find CPR bacteria to be unique to CG-1 as it is closer to the geyser borehole and more frequently exposed to geyser discharge during eruptions (Fig. 1A-B).

To confirm that differences in unique bins between CG-1 and CG-10 are not due to sequencing depths, we compared mappable reads. CG-1 had 97.5% mappable reads (averaging 219,594,576 reads between both sequencing facilities), while CG-10 had 93% (204,434,731 reads). Although CG-10 has fewer mappable reads, significantly more unique bins were recovered, suggesting higher diversity at CG-10 compared to CG-1 (Table S6). In summary, our analysis demonstrated differences in community composition between the two sites suggesting potential differences in ecological processes that shape microbial community compositions between the two sites.

### *Rhodobacterales* are most abundant across sampling sites suggesting functional importance in travertine communities

To further assess community structures, we calculated the relative abundances of each bin based on metagenomic read counts, which can be a measure for cell/genome copy numbers, as well as metaproteomics-based biomass quantification (42). We found that *Rhodobacterales* were most abundant in both communities and based on both methods (Fig. 3, Supplementary File 2). At CG-1, *Rhodobacterales* contributed a total of 14.5% of metagenomic reads and 27.3% of proteinaceous biomass. At CG-10, *Rhodobacterales* were equally abundant across both methods with 29.1% of metagenomic reads and 29.2% of proteinaceous biomass. Among *Rhodobacterales*, the most prevalent genera included *Roseovarius*, *Roseicyclus*, *Rubrimonas*, *Fluviibacterium*, and *Oceaniglobus*, observed in both sites and across both methods. Notably, at CG-10, *Tateyamaria*, *Pseudorhodobacter*, *Roseinatrobacter*, and *Nioella* were also highly abundant (Supplementary File 2). While the majority of these genera are characterized as strictly aerobic chemoheterotrophs, *Rubrimonas* and *Roseicyclus* are aerobic anoxygenic phototrophs (AAP) (53). Our findings revealed a significant difference in the abundance of *Rubrimonas* between the CG-1 and CG-10 environments. At CG-1, *Rubrimonas* was the most abundant genus within *Rhodobacterales*, as shown by both our metagenomics and metaproteomics data. However, at CG-10, this was only evident in the metagenomics data. Instead, at CG-10, *Roseovarius* was the most abundant genus in terms of proteinaceous biomass (Supplementary File 2).

**Fig. 3:**
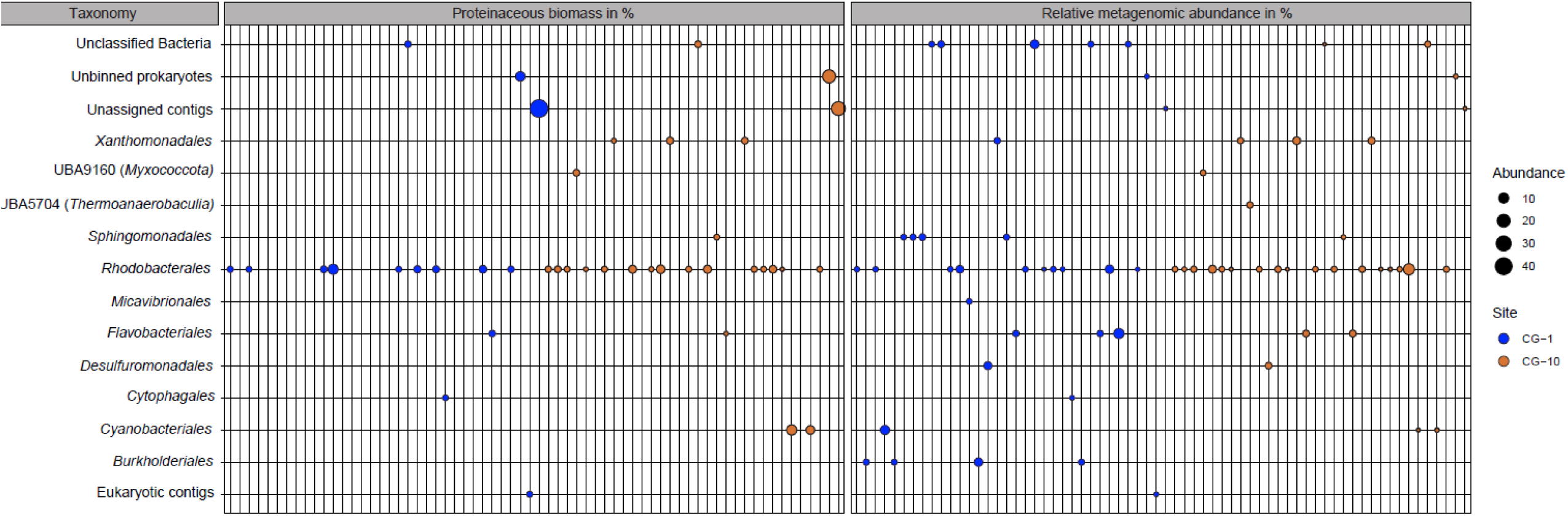
Taxonomy, metagenomics-based, and metaproteomics-based abundance of taxa (bins) in microbial communities recovered from CG travertine. Each bubble represents one bin/organism. Only organisms contributing more than 1% of relative bin abundance based on metagenomic read mapping and/or metaproteomics-based proteinaceous biomass assessment are displayed. Proteinaceous biomass was averaged across three replicates respectively. Empty columns in the proteinaceous biomass panel are organisms that passed the abundance threshold for read-based bin abundance but were not detected at all in the proteinaceous biomass calculation. Unassigned bacteria represent bins that could not be classified with GTDB-Tk. Unbinned prokaryotes are prokaryotic contigs that were classified as such by Whokaryote but could not be assigned to a bin by MetaBat2. Unassigned contigs are short contigs that were filtered out by Whokaryote and were manually recovered.

*Rhodobacterales* are capable of a diversity of metabolisms but mainly comprise aerobic photo-or chemoheterotrophs, with some also being capable of autotrophy and phototropy in anaerobic environments (53). We found abundant proteins for phototrophy, uptake of organic substrates, and autotrophic carbon fixation in *Rhodobacterales* bins, often together in the same bin indicating a mixotrophic lifestyle (Supplementary File 3 & 4). The high abundance of mixotrophic microorganisms could indicate nutrient limitation, as it has been shown previously that mixotrophy is favored in oligotrophic/nutrient-limited environments (54,55).

Despite their low abundance based on metagenomic reads, unassigned contigs contributed significantly to the proteinaceous biomass at CG-1 and CG-10, accounting for 44.54% and 22.71% of the total proteinaceous biomass, respectively. We also found that *Flavobacteriales, Burkolderiales,* and *Cyanobacteriales* were highly abundant at CG-1 based on metagenomic reads (13.5%, 9.6%, and 7.3% respectively) and low abundant or absent in our metaproteomics data (Fig. 3, Supplementary File 2). In contrast, we found *Cyanobacteriales* to be low abundant in metagenomic data of CG-10 (0.3%) but contributed 15.1% of proteinaceous biomass at this site. In summary, we found various genera of *Rhodobacterales* were most abundant in terms of metagenomic read number and proteinaceous biomass suggesting that they play important functional roles in CG communities.

While *Rubrimonas* and *Roseicyclus* have not been directly associated with MICP thus far, there has been some exploration of anoxygenic photosynthetic bacteria in MICP research. However, despite experimental evidence, the mechanisms by which these bacteria facilitate CaCO_3_ precipitation remain to be fully understood (56,57). Similar to CG communities, *Alphaproteobacteria*, to which the *Rhodobacterales* belong, and *Cyanobacteria* are commonly found to co-exist in stromatolites (58,59) suggesting that similar community structures might be at play in CG travertine.

### Formation of Crystal Geyser travertine is likely mediated by microbial photosynthetic activity

Several microbial physiologies have been linked to MICP, including photosynthesis, anaerobic methane oxidation, denitrification, urea hydrolysis, ammonification, and sulfate reduction. We screened our metaproteomic datasets for key enzymes involved in these processes, such as RuBisCO, photosystem proteins, methyl-coenzyme M reductases, nitrate/nitrite/nitric oxide/nitrous oxide reductases, urease, and sulfite reductases. We identified 5,632 proteins in travertine from CG-1 and 1,146 proteins in CG-10 (Supplementary File 3 and 4). Of these, 57 proteins at CG-1 and 24 at CG-10 were of interest (Supplementary File 5). We found proteins for photosynthesis and incomplete denitrification, but none for methane oxidation, and only one each for sulfate reduction and urea hydrolysis.

### Photosynthesis

At CG-1, we identified 41 proteins involved in photosynthetic activity with the most abundant being a RuBisCO Form I large chain belonging to *Pikeienuella sp*. (*Rhodobacterales*) which contributed to 0.2% of all identified proteins in terms of abundance at this site. At CG-10, we exclusively identified photosynthesis-related proteins among our proteins of interest with the most abundant being a photosystem I subunit belonging to *Cyanobacteriales* of the genus *Planktothrix sp.* (Supplementary File 5). This protein was not only abundant amongst our proteins of interest but the second most abundant protein at this site contributing almost 1.5% of all detected proteins. Other proteins related to photosynthesis included several photosystem I and II subunits as well as photosynthetic reaction center subunits detected at both sites. Most of the photosynthetic proteins at CG-1 belonged to several genera of *Cyanobacteriales* (*Geminocystis sp*., and *Planktothrix* sp.) as well as *Rhodobacterales* (mainly *Rubrimonas* and *Fluviibacterium*) indicating that oxygenic photosynthesis and potentially anoxygenic photosynthesis could play a role at CG-1. Unfortunately, we could not taxonomically assign most photosynthesis-related proteins at CG-10 and only 3 out of 24 proteins were linked to *Cyanobacteriales* and one to *Rhodobacterales* while the remaining proteins were derived from unassigned contigs. Given the large contribution of proteinaceous biomass of *Cyanobacteriales* and the fact that other identified proteins at this site hint towards oxygenic photosynthesis (e.g. highly abundant carboxysome proteins), we expect that *Cyanobacteriales* and thus oxygenic photosynthesis play a more important role at CG-10 as compared to CG-1. MICP via oxygenic photosynthesis has been extensively studied as it contributes to many carbonate formation processes such as reefs, lacustrine whitings, carbonated sediments, and stromatolites (60). In most of these settings, MICP is driven by *Cyanobacteria* and has been linked to their Carbon Concentration Mechanism (CCM). The CCM functions to concentrate CO_2_ intracellularly, optimizing RuBisCO’s efficiency for carbon fixation. In brief, HCO_3_^−^ enters the cell via transporters and is then transported to carboxysomes — a protein-enclosed micro-compartment containing both carbonic anhydrase and RuBisCO. Within the carboxysome, carbonic anhydrase converts HCO_3_^−^ into CO_2_, which is then fixed into sugars by RuBisCO. This transformation consumes H^+^, resulting in an elevated pH outside the cell. The ensuing alkalization, in conjunction with Ca^2+^ accumulation at the cell surface, amplifies the CaCO_3_ saturation state in the surrounding solution, likely leading to CaCO_3_ precipitation (Fig. 4).

**Fig. 4.**
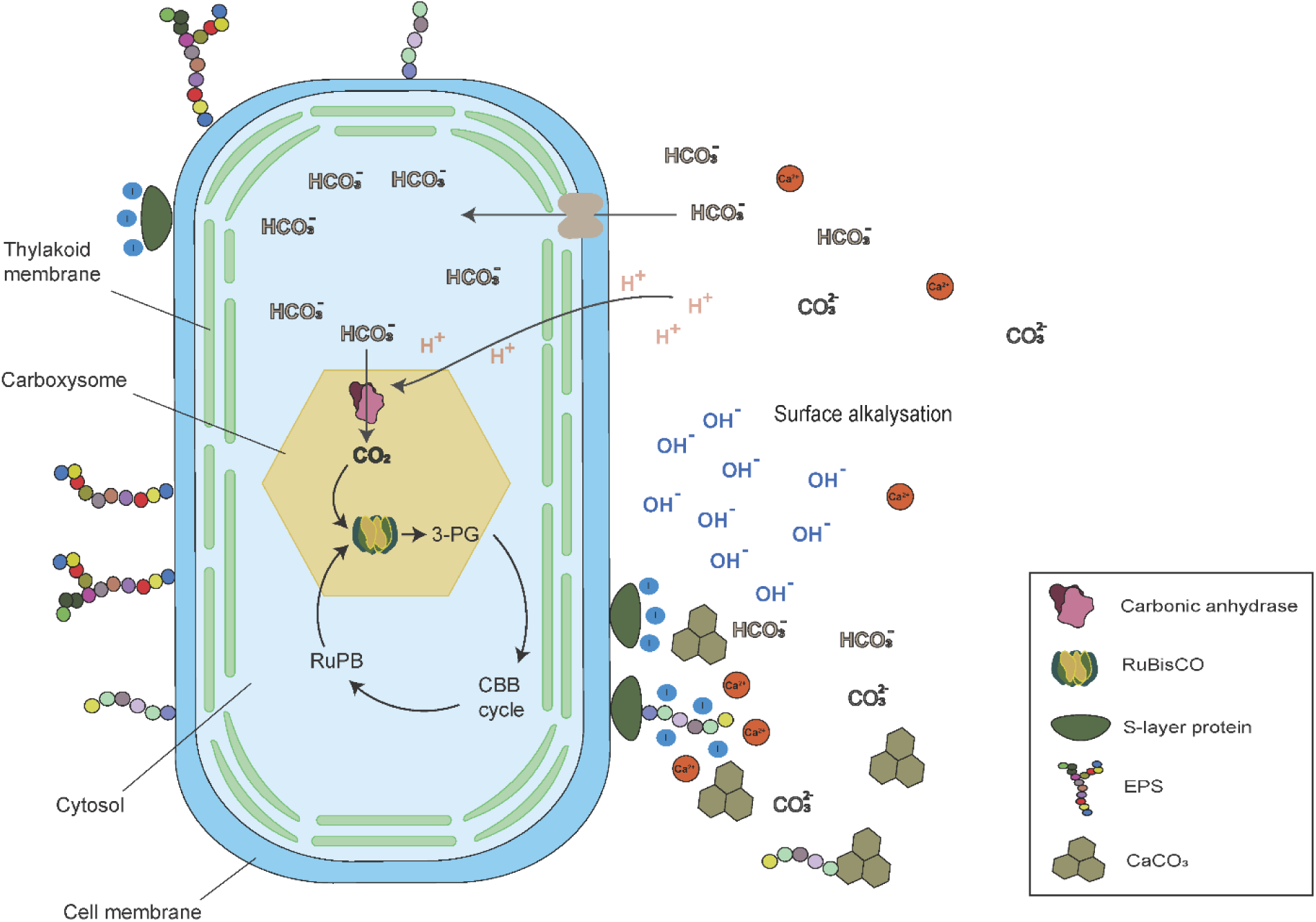
Schematic depiction of the CO_2_-concentrating mechanism (CCM) linked to MICP in a cyanobacterial cell. Modified from Kamennaya et al., 2012 (10).

To further confirm the photosynthetic influence on carbonate formation at CG, we measured stable carbon isotope ratios of travertine samples collected from CG-1 and CG-10. Photosynthetic organisms, particularly ones that use RuBisCO for carbon fixation, preferentially use ^12^C leading to a characteristic enrichment of ^13^C in their extracellular environment (61). This ^13^C enrichment leads to microbially induced carbonates being isotopically “heavier” as compared to carbonates produced solely by abiotic processes. Our carbon isotope analysis of CG travertine revealed a strong ^13^C enrichment consistent with the role of microbial photosynthesis in travertine formation at CG. The average δ^13^C value measured at CG-1 was +7.32‰±0.03‰ and +7.38‰±‰ 0.15 at CG-10. These values are notably higher than previously reported δ^13^C_DIC_ derived from the geyser discharge which was 5.0‰±1.4‰ (Fig. 5C, Table S7) (51). We acknowledge that CO_2_ degassing could also be a likely source of ^13^C enrichment at CG, given the CO_2_-enriched water from the geyser and the climate at CG. If CO_2_ degassing were to be the main driver of carbonate precipitation, we would expect that to be reflected in the isotopic composition, as this would lead to a characteristic enrichment of both δ ^13^C and δ^18^O (62,63). However, the δ^18^O composition of the CG carbonates matched the annual meteoric δ^18^O of the Green River area and the average water temperature (64,65). Since microbial photosynthetic activity does not affect δ^18^O in carbonates (66), the absence of δ^18^O enrichment from CO_2_ degassing confirms that our carbonate data reflects the primary precipitation conditions in the CG environment (Table S7).

**Fig 5:**
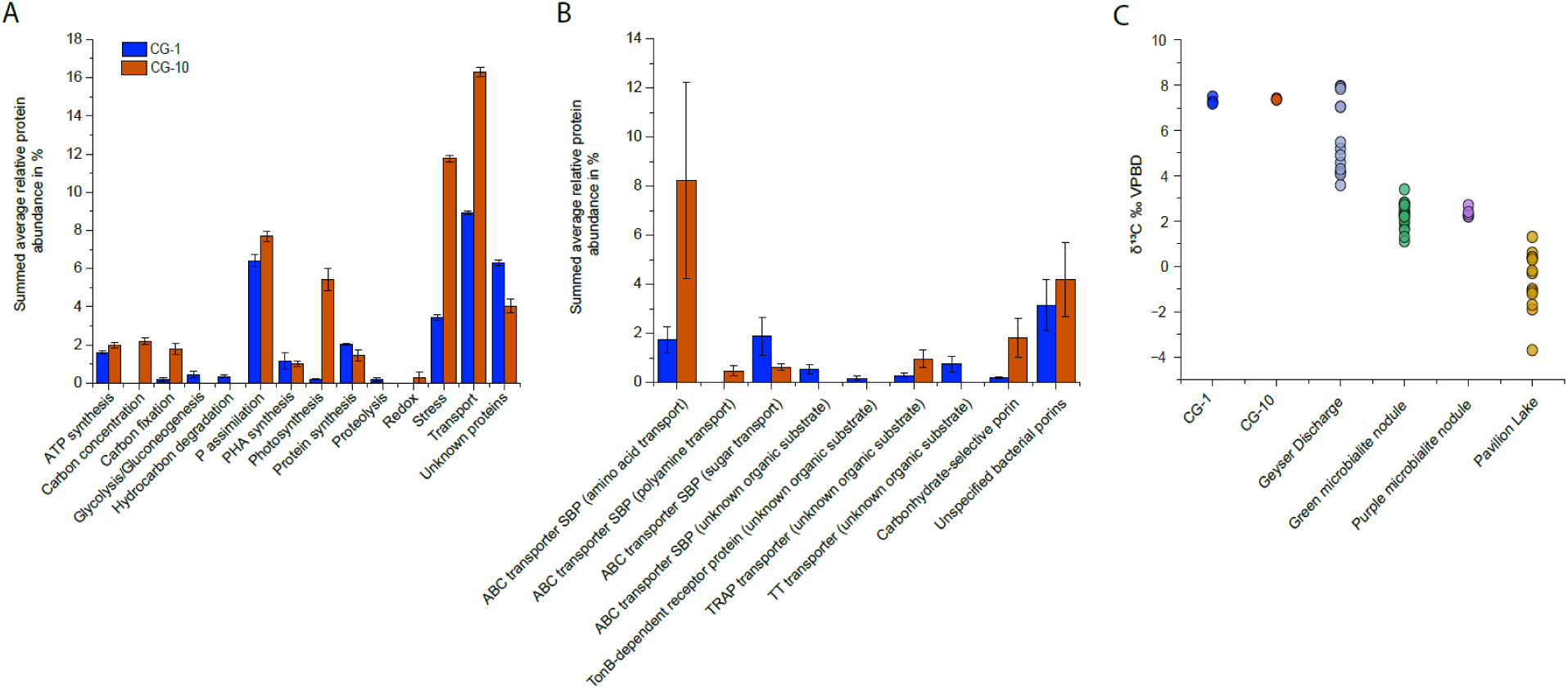
Abundance trends of detected proteins in CG travertine and inference of MICP mechanism. A) Overview of the metabolic categories of the 100 most abundant proteins, B) abundances of proteins for the uptake of specific organic substrates, C) stable carbon isotope analysis of CG travertine and water (65) and comparison samples from microbialite nodules from Pavilion Lake, B.C. Canada (72). Relative protein abundances are averages across three replicates respectively. Averages were summed up per category to show the average contribution of each specific metabolic category at each site respectively. Error bars represent average standard deviation from the mean within each category. SBP: Substrate-binding protein.

To further put this in perspective, we compared our measured δ^13^C values to previously reported δ^13^C values of micro-stromatolitic nodules from freshwater microbialites located in Pavilion Lake, B.C. Canada (61) (Fig. 5C). These nodules were associated with *Cyanobacteria* and thus expected to carry similar photosynthetic biosignatures. Indeed, the authors found strong evidence for photosynthetic influence on precipitation within the nodule microenvironments with an average carbonate δ^13^C value of +2.3‰ (± 0.54) in green nodules and +2.36‰ (± 0.21) in purple nodules. Interestingly, the authors observed a δ^13^C enrichment of ∼2‰ when compared to equilibrium, which is a similar offset to what we observed in CG travertine. It is noteworthy, however, that the geyser water was notably enriched in ^13^C which is thought to be due to the complex microbial communities that reside in the deep surface of the aquifer which have been proposed to perform carbon fixation thus leading to this characteristic enrichment of ^13^C in the geyser discharge (61, 65, 66). Nevertheless, the reported average δ^13^C_DIC_ of 5.0‰±1.4‰ is still considerably lower than the measured carbonate δ^13^C values, even when taking the expected δ^13^C enrichment during precipitation into account, thus further supporting our hypothesis that travertine formation at CG is likely mediated by microbial photosynthetic activity.

### Denitrification

We detected several enzymes involved in microbial denitrification in our metaproteomic data from CG-1 (Supplementary File 5) including the catalytic subunits of 3 respiratory nitrate reductases and 8 nitrite reductases. All the detected enzymes either belonged to *Rhodobacterales* or were derived from short, unassigned contigs. We did not identify any other enzymes associated with denitrification, such as nitric oxide reductase or nitrous oxide reductase. This absence could be attributed either to the absence of complete denitrifiers in CG communities or to the fact that most of the enzymes involved are membrane-bound and thus harder to detect by LC-MS/MS based metaproteomics. We also measured bulk δ^15^N of CG travertine and did not find the characteristic enrichment of ^15^N that indicates microbial denitrification thus suggesting that denitrification might play a minor role in CG sediments (67) (Table S3).

Denitrification is a ubiquitous process in nature involving a series of enzyme-catalyzed reduction steps reducing nitrate to nitrogen gas (N_2_). If denitrifying microorganisms use an organic substrate as an electron donor they generate dissolved inorganic carbon (DIC), as well as consume H^+^ through the nitrate reduction reactions leading to an increase in local pH. In an aqueous solution, the generated DIC will dissociate into CO_2_, HCO_3_^−^ and CO_3_^2-^ and if the pH is high and excess Ca^2+^ is present, CaCO_3_ will precipitate. Our findings indicate that denitrification might not serve as a predominant MICP mechanism at CG, especially in the sampled aerobic zone of the travertine. Nevertheless, it could still play a minor role alongside photosynthesis.

### Urea hydrolysis

We detected one catalytic subunit of urease at CG-1, which was the lowest abundance protein among our selected proteins of interest (Supplementary File 5). Although the metaproteomics data did not show the presence of other urease subunits, we identified the necessary remaining subunits and accessory genes in the gene neighborhood of the detected protein, confirming that this bacterium has the metabolic potential for urea hydrolysis. Our results suggest that while urea hydrolysis is likely not a dominant MICP mechanism at CG it may play a minor role in conjunction with photosynthesis and nitrate reduction.

### High abundance of high-affinity phosphate uptake proteins indicates that community is phosphate limited

Multiple community members expressed proteins for high-affinity phosphate uptake at high abundances (Fig. 5A, Supplementary File 3 & 4). The periplasmic substrate-binding protein (SPB) PstS of the phosphate transport system was the most abundant protein at both sites contributing 1.04 % of all identified proteins at CG-1 and 2.3% at CG-10. While some of the subunits for the transporters were not detected in the metaproteomes likely due to their lower copy number and embedding in the membrane (68), we did find the remaining genes of the *pstSCAB* operon in the genomic neighborhood of the abundant PstS proteins. The *pstSCAB* operon encoded high-affinity transporter is an important mechanism by which bacteria can adapt to phosphate-deficient environments and acquire this essential nutrient for their growth and survival. Under low phosphate conditions, the expression of *pstSCAB* is stimulated, resulting in an enhanced ability of bacteria to take up phosphate even at extremely low concentrations (69). To further test the hypothesis that the microbial communities at CG are P limited, we investigated the presence of other proteins of the Pho regulon, which includes genes involved in bacterial P management including the *pstSCAB* operon. We identified alkaline phosphatase, which is part of the Pho regulon, at both CG-1 and CG-10 (Supplementary File 3 & 4). At CG-1, our analysis revealed the presence of two distinct alkaline phosphatases, contributing 0.16% and 0.0003%, respectively, to the overall protein abundance across all identified proteins at this site. In contrast, in CG-10 samples we identified a single alkaline phosphatase with an abundance of 0.04%. The expression of alkaline phosphatase in addition to the high PstS abundances further supports the hypothesis that microbiomes at CG are P limited.

To further support our metaproteomics-based prediction of phosphate limitation, we carefully analyzed our SEM-EDX elemental data of CG travertine (Table S1-2). Our findings revealed that phosphorus concentrations were below the detectable threshold in all samples.

An additional line of evidence that points to nutrient limitation in the CG travertine microbial communities is the abundant expression of proteins involved in the production of the storage compound synthesis of polyhydroxyalkanoates (PHAs). We found Acetoacetyl-CoA reductases (PhaB) among the top 100 most abundant proteins, which are involved PHA synthesis (Supplementary File 6). PHAs are intracellular energy and carbon storage granules that are commonly produced by bacteria in response to nutrient limitations, such as low levels of nitrogen and phosphorus. By synthesizing PHAs, bacteria can store excess carbon and energy in the form intracellular granules, which can be used later when nutrients become available or carbon and energy become scarce (70). Overall, our results suggest that CG travertine is a phosphate-limited environment and that P-limitation shapes the microbial communities and their physiology in this ecosystem.

### High abundance of ABC transporters suggests active uptake of various organic substrates

We identified multiple community members that expressed highly abundant proteins for the uptake of various organic substrates (Fig. 5B). Transport-related proteins made up a third of the abundance of the top 100 most abundant proteins across all three replicates at both sites (Fig. 5A-B). At CG-1, the most abundant transport-related proteins were unclassified bacterial porins and ABC transporter substrate-binding proteins (SBPs) for sugars and amino acids. Similarly, ABC transporter SBPs for the uptake of amino acids were the most abundant transport-related proteins at CG-10 followed by unspecified bacterial porins and carbohydrate specific porins (Fig. 5B). ABC transporters are found in all domains of life and are involved in the translocation of various substrates across cellular membranes. In contrast to other transporters, ABC transporters have a high energy requirement as two ATP molecules are hydrolyzed per substrate molecule transported (71) and thus are usually only used when substrate concentrations are too low to enable transport by other mechanisms. The detection of high-affinity substrate-binding proteins for amino acids, sugars, and carbohydrates at CG suggests the presence of these substrates, but their concentration range is likely low (72–74). The high abundance of ABC transporter proteins may additionally indicate nutrient scarcity, the abundant SBPs for amino acids, for instance, may indicate that the communities are limited in nitrogen. Nitrogen is an essential component for amino acid synthesis, and the high abundance of transport proteins may suggest that the communities are actively scavenging for amino acids to meet their nitrogen requirements.

## Conclusion and Future Directions

We conducted a metagenomic-metaproteomic study coupled with stable carbon isotope analyses to unravel potential mechanisms of MICP at a site where the microbial community is involved in travertine formation. Our study highlights that photosynthesis carried out by *Cyanobacteria* and potentially *Rubrimonas* and *Roseicyclus* could be a driver of MICP in terrestrial environments, in addition to its role in marine stromatolites and whiting events. Additionally, in contrast to most research on MICP, which has been conducted using pure cultures of bacteria under controlled laboratory conditions with high nutrient availability, we found that MICP occurs in natural environments at CG despite nutrient limitation (P, N), providing new insights into constraints of MICP under environmental conditions. As of right now, we do not know if nutrient limitation at CG limits MICP or if nutrient limitation is promoting MICP. This could be tested in future field experiments with nutrient addition. Knowing how nutrient supply impacts MICP would be critical for the development of MICP-based engineering solutions as high costs associated with microbial substrates and calcifying media are one of the main challenges for the widespread adoption of MICP. Nutrients, such as nitrogen, and phosphorus are essential for the growth of bacteria and the availability and concentration of these nutrients in the environment may significantly affect the rate and efficiency of MICP.

While our results provide valuable insights, they are limited by the lack of information about the rate at which MICP occurs at CG and which community members are actively contributing to MICP at CG. Future investigations could involve the isolation of specific photosynthetic community members that are found in CG travertine or the use of close relatives from culture collections. By conducting laboratory experiments to test their MICP capabilities, the rate of precipitation could be measured. Additionally, rate of carbonate precipitation could also be measured with radiotracers of ^45^Ca in the field. These techniques would enable the determination of the efficiency of the specific photosynthetic community members in facilitating MICP, and provide further insight into the underlying mechanisms of MICP.

It is important to acknowledge that photosynthetically-produced CaCO_3_ comes with certain limitations as it can only be used in aboveground applications, such as building and construction materials or, depending on the light penetration depths and porosity of the soil matrix, near ground-surface and stabilization applications (75). Addressing these issues will be critical for the development of reliable and effective bio-mediated technologies for soil stabilization and construction, and will require interdisciplinary collaborations between microbiologists, geologists, materials scientists, and engineers.

## Acknowledgments

We thank Marlee Reed for teaching us the carbonate quantification method and Harald Gruber-Vodicka and Mike Lee for bioinformatics support and fruitful discussions. This work was performed in part at the Analytical Instrumentation Facility (AIF) at North Carolina State University, which is supported by the State of North Carolina and the National Science Foundation (award number ECCS-2025064). The AIF is a member of the North Carolina Research Triangle Nanotechnology Network (RTNN), a site in the National Nanotechnology Coordinated Infrastructure (NNCI).

## Contributions

MV performed all data analysis, research, and writing. EH, AG and LB managed and assisted sample collection and field documentation. EH performed geochemical analyses. BM conceptually designed experiments. MK guided all research efforts, including analyses and writing. All authors read and approved the final manuscript.

## Ethics declaration

The authors declare no competing interests.

## References

1. Hans Wedepohl K. The composition of the continental crust. Geochimica et Cosmochimica Acta. 1995 Apr;59(7):1217–32.

2. Gadd GM. Metals, minerals and microbes: geomicrobiology and bioremediation. Microbiology. 2010 Mar 1;156(3):609–43.

3. Mortensen BM, Haber MJ, DeJong JT, Caslake LF, Nelson DC. Effects of environmental factors on microbial induced calcium carbonate precipitation: Environmental factors on MICP. Journal of Applied Microbiology. 2011 Aug;111(2):338–49.

4. Dupraz C, Visscher PT, Baumgartner LK, Reid RP. Microbe-mineral interactions: early carbonate precipitation in a hypersaline lake (Eleuthera Island, Bahamas): Microbe-mineral interactions, Eleuthera Island, Bahamas. Sedimentology. 2004 Jun 17;51(4):745–65.

5. Reeburgh WS. Oceanic Methane Biogeochemistry. Chem Rev. 2007 Feb 1;107(2):486–513.

6. van Paassen LA, Daza CM, Staal M, Sorokin DY, van der Zon W, van Loosdrecht MarkCM. Potential soil reinforcement by biological denitrification. Ecological Engineering. 2010 Feb;36(2):168–75.

7. Reyes C, Schneider D, Lipka M, Thürmer A, Böttcher ME, Friedrich MW. Nitrogen Metabolism Genes from Temperate Marine Sediments. Mar Biotechnol. 2017 Apr;19(2):175–90.

8. Gallagher KL, Kading TJ, Braissant O, Dupraz C, Visscher PT. Inside the alkalinity engine: the role of electron donors in the organomineralization potential of sulfate-reducing bacteria. Geobiology. 2012 Nov;10(6):518–30.

9. Giuffre AJ, Hamm LM, Han N, De Yoreo JJ, Dove PM. Polysaccharide chemistry regulates kinetics of calcite nucleation through competition of interfacial energies. Proc Natl Acad Sci USA. 2013 Jun 4;110(23):9261–6.

10. Kamennaya N, Ajo-Franklin C, Northen T, Jansson C. Cyanobacteria as Biocatalysts for Carbonate Mineralization. Minerals. 2012 Oct 29;2(4):338–64.

11. Visscher PT, Stolz JF. Microbial mats as bioreactors: populations, processes, and products. Palaeogeography, Palaeoclimatology, Palaeoecology. 2005 Apr;219(1–2):87–100.

12. Lal R. Carbon sequestration. Phil Trans R Soc B. 2008 Feb 27;363(1492):815–30.

13. Liu P, Zhang Y, Tang Q, Shi S. Bioremediation of metal-contaminated soils by microbially-induced carbonate precipitation and its effects on ecotoxicity and long-term stability. Biochemical Engineering Journal. 2021 Feb;166:107856.

14. Karimi N, Mostofinejad D. Bacillus subtilis bacteria used in fiber reinforced concrete and their effects on concrete penetrability. Construction and Building Materials. 2020 Jan;230:117051.

15. Wu Y, Li H, Li Y. Biomineralization Induced by Cells of Sporosarcina pasteurii: Mechanisms, Applications and Challenges. Microorganisms. 2021 Nov 21;9(11):2396.

16. Torres-Aravena Á, Duarte-Nass C, Azócar L, Mella-Herrera R, Rivas M, Jeison D. Can Microbially Induced Calcite Precipitation (MICP) through a Ureolytic Pathway Be Successfully Applied for Removing Heavy Metals from Wastewaters? Crystals. 2018 Nov 21;8(11):438.

17. Barth JA, Chafetz HS. Cool water geyser travertine: Crystal Geyser, Utah, USA. Pufahl P, editor. Sedimentology. 2015 Apr;62(3):607–20.

18. Cosmidis J, O’Reilly S, Ellison E, Crispin K, Diercks D, Templeton A. Carbonate polymorphism controlled by microbial iron redox dynamics at a natural CO2 leakage site (Crystal Geyser, Utah) [Internet]. Geochemistry; 2021 Jul [cited 2023 Apr 21]. Available from: http://eartharxiv.org/repository/view/2523/

19. Takashima C, Okumura T, Nishida S, Shimamoto T, Koike H, Kano A. Microbial Control on Lamina Formation in a Travertine of Crystal Geyser, Utah. In: Advances in Stromatolite Geobiology [Internet]. Berlin, Heidelberg: Springer Berlin Heidelberg; 2011 [cited 2023 Apr 21]. p. 123–33. (Lecture Notes in Earth Sciences; vol. 131). Available from: http://link.springer.com/10.1007/978-3-642-10415-2_7

20. Menke S, Gillingham MAF, Wilhelm K, Sommer S. Home-Made Cost Effective Preservation Buffer Is a Better Alternative to Commercial Preservation Methods for Microbiome Research. Front Microbiol [Internet]. 2017 Jan 31 [cited 2023 Apr 21];8. Available from: http://journal.frontiersin.org/article/10.3389/fmicb.2017.00102/full

21. Jensen M, Wippler J, Kleiner M. Evaluation of RNA *later* as a Field-Compatible Preservation Method for Metaproteomic Analyses of Bacterium-Animal Symbioses. Gralnick JA, editor. Microbiol Spectr. 2021 Oct 31;9(2):e01429–21.

22. Gray MA, Pratte ZA, Kellogg CA. Comparison of DNA preservation methods for environmental bacterial community samples. FEMS Microbiol Ecol. 2013 Feb;83(2):468–77.

23. O’Toole C, Liu Q, Montoya BM, Kananizadeh N, Odle W. The Effect of Microbial Induced Carbonate Precipitation on Fine-Grained Mine Tailings. In: Geo-Congress 2022 [Internet]. Charlotte, North Carolina: American Society of Civil Engineers; 2022 [cited 2023 Apr 21]. p. 335–46. Available from: http://ascelibrary.org/doi/10.1061/9780784484012.035

24. Andrews S. FASTQC. A quality control tool for high throughput sequence data. 2010.

25. Bushnell B. BBMap: A Fast, Accurate, Splice-Aware Aligner. 2014 Mar; Available from: https://www.osti.gov/biblio/1241166

26. Nurk S, Meleshko D, Korobeynikov A, Pevzner PA. metaSPAdes: a new versatile metagenomic assembler. Genome Res. 2017 May;27(5):824–34.

27. Gruber-Vodicka HR, Seah BKB, Pruesse E. phyloFlash: Rapid Small-Subunit rRNA Profiling and Targeted Assembly from Metagenomes. Arumugam M, editor. mSystems. 2020 Oct 27;5(5):e00920–20.

28. Li D, Liu CM, Luo R, Sadakane K, Lam TW. MEGAHIT: an ultra-fast single-node solution for large and complex metagenomics assembly via succinct *de Bruijn* graph. Bioinformatics. 2015 May 15;31(10):1674–6.

29. Pronk LJU, Medema MH. Whokaryote: distinguishing eukaryotic and prokaryotic contigs in metagenomes based on gene structure. Microbial Genomics [Internet]. 2022 May 3 [cited 2023 Apr 21];8(5). Available from: https://www.microbiologyresearch.org/content/journal/mgen/10.1099/mgen.0.000823

30. Hyatt D, Chen GL, LoCascio PF, Land ML, Larimer FW, Hauser LJ. Prodigal: prokaryotic gene recognition and translation initiation site identification. BMC Bioinformatics. 2010 Dec;11(1):119.

31. Chaumeil PA, Mussig AJ, Hugenholtz P, Parks DH. GTDB-Tk: a toolkit to classify genomes with the Genome Taxonomy Database. Hancock J, editor. Bioinformatics. 2019 Nov 15;btz848.

32. Bowers RM, Kyrpides NC, Stepanauskas R, Harmon-Smith M, Doud D, Reddy TBK, et al. Minimum information about a single amplified genome (MISAG) and a metagenome-assembled genome (MIMAG) of bacteria and archaea. Nature Biotechnology. 2017 Aug 1;35(8):725–31.

33. Lee MD. GToTree: a user-friendly workflow for phylogenomics. Ponty Y, editor. Bioinformatics. 2019 Oct 15;35(20):4162–4.

34. Hug LA, Baker BJ, Anantharaman K, Brown CT, Probst AJ, Castelle CJ, et al. A new view of the tree of life. Nat Microbiol. 2016 Apr 11;1(5):16048.

35. Eddy SR. Accelerated Profile HMM Searches. Pearson WR, editor. PLoS Comput Biol. 2011 Oct 20;7(10):e1002195.

36. Edgar RC. High-accuracy alignment ensembles enable unbiased assessments of sequence homology and phylogeny [Internet]. Bioinformatics; 2021 Jun [cited 2023 Apr 21]. Available from: http://biorxiv.org/lookup/doi/10.1101/2021.06.20.449169

37. Capella-Gutiérrez S, Silla-Martínez JM, Gabaldón T. trimAl: a tool for automated alignment trimming in large-scale phylogenetic analyses. Bioinformatics. 2009 Aug 1;25(15):1972–3.

38. Price MN, Dehal PS, Arkin AP. FastTree 2 – Approximately Maximum-Likelihood Trees for Large Alignments. Poon AFY, editor. PLoS ONE. 2010 Mar 10;5(3):e9490.

39. Letunic I, Bork P. Interactive Tree Of Life (iTOL) v5: an online tool for phylogenetic tree display and annotation. Nucleic Acids Research. 2021 Jul 2;49(W1):W293–6.

40. Qian C, Hettich RL. Optimized Extraction Method To Remove Humic Acid Interferences from Soil Samples Prior to Microbial Proteome Measurements. J Proteome Res. 2017 Jul 7;16(7):2537–46.

41. Wiśniewski JR, Zougman A, Nagaraj N, Mann M. Universal sample preparation method for proteome analysis. Nat Methods. 2009 May;6(5):359–62.

42. Kleiner M, Thorson E, Sharp CE, Dong X, Liu D, Li C, et al. Assessing species biomass contributions in microbial communities via metaproteomics. Nat Commun. 2017 Nov 16;8(1):1558.

43. Perez-Riverol Y, Bai J, Bandla C, García-Seisdedos D, Hewapathirana S, Kamatchinathan S, et al. The PRIDE database resources in 2022: a hub for mass spectrometry-based proteomics evidences. Nucleic Acids Research. 2022 Jan 7;50(D1):D543–52.

44. Heath JE, Lachmar TE, Evans JP, Kolesar PT, Williams AP. Hydrogeochemical characterization of leaking, carbon dioxide-charged fault zones in east-central Utah, with implications for geologic carbon storage. In: McPherson BJ, Sundquist ET, editors. Geophysical Monograph Series [Internet]. Washington, D. C.: American Geophysical Union; 2009 [cited 2023 Apr 21]. p. 147–58. Available from: https://onlinelibrary.wiley.com/doi/10.1029/2006GM000407

45. Probst AJ, Ladd B, Jarett JK, Geller-McGrath DE, Sieber CMK, Emerson JB, et al. Differential depth distribution of microbial function and putative symbionts through sediment-hosted aquifers in the deep terrestrial subsurface. Nat Microbiol. 2018 Jan 29;3(3):328–36.

46. Lenher V, Merrill HB. THE SOLUBILITY OF SILICA. J Am Chem Soc. 1917 Dec;39(12):2630–8.

47. Podosokorskaya OA, Kadnikov VV, Gavrilov SN, Mardanov AV, Merkel AY, Karnachuk OV, et al. Characterization of Melioribacter roseus gen. nov., sp. nov., a novel facultatively anaerobic thermophilic cellulolytic bacterium from the class Ignavibacteria, and a proposal of a novel bacterial phylum Ignavibacteriae. Environmental Microbiology. 2013;15(6):1759–71.

48. Cavalier-Smith T, Chao EEY. Multidomain ribosomal protein trees and the planctobacterial origin of neomura (eukaryotes, archaebacteria). Protoplasma. 2020 May 1;257(3):621–753.

49. Ward LM, Lingappa UF, Grotzinger JP, Fischer WW. Microbial mats in the Turks and Caicos Islands reveal diversity and evolution of phototrophy in the Chloroflexota order Aggregatilineales. Environmental Microbiome. 2020 Dec;15(1):9.

50. Nakahara N, Nobu MK, Takaki Y, Miyazaki M, Tasumi E, Sakai S, et al. Aggregatilinea lenta gen. nov., sp. nov., a slow-growing, facultatively anaerobic bacterium isolated from subseafloor sediment, and proposal of the new order Aggregatilineales ord. nov. within the class Anaerolineae of the phylum Chloroflexi. International Journal of Systematic and Evolutionary Microbiology. 2019 Apr 1;69(4):1185–94.

51. Probst AJ, Elling FJ, Castelle CJ, Zhu Q, Elvert M, Birarda G, et al. Lipid analysis of CO2-rich subsurface aquifers suggests an autotrophy-based deep biosphere with lysolipids enriched in CPR bacteria. ISME J. 2020 Jun;14(6):1547–60.

52. Emerson JB, Thomas BC, Alvarez W, Banfield JF. Metagenomic analysis of a high carbon dioxide subsurface microbial community populated by chemolithoautotrophs and bacteria and archaea from candidate phyla: High CO _2_ subsurface metagenomics. Environ Microbiol. 2016 Jun;18(6):1686–703.

53. DeLong EF, Rosenberg E, Lory S, Stackebrandt E, Thompson F, editors. The Prokaryotes. Alphaproteobacteria and betaproteobacteria / Eugene Rosenberg (editor-in-chief) ; Edward F. DeLong, Stephen Lory, Erko Stackebrandt and Fabiano Thompson (eds.). Fourth edition. Heidelberg New York Dordrecht London: Springer Reference; 2014. 1012 p.

54. Thingstad TF, Havskum H, Garde K, Riemann B. On the Strategy of “Eating Your Competitor”: A Mathematical Analysis of Algal Mixotrophy. Ecology. 1996 Oct;77(7):2108–18.

55. Edwards KF. Mixotrophy in nanoflagellates across environmental gradients in the ocean. Proc Natl Acad Sci USA. 2019 Mar 26;116(13):6211–20.

56. Bosak T, Greene SE, Newman DK. A likely role for anoxygenic photosynthetic microbes in the formation of ancient stromatolites. Geobiology. 2007 Jun;5(2):119–26.

57. Bundeleva IA, Shirokova LS, Bénézeth P, Pokrovsky OS, Kompantseva EI, Balor S. Calcium carbonate precipitation by anoxygenic phototrophic bacteria. Chemical Geology. 2012 Jan;291:116–31.

58. Baumgartner LK, Spear JR, Buckley DH, Pace NR, Reid RP, Dupraz C, et al. Microbial diversity in modern marine stromatolites, Highborne Cay, Bahamas. Environmental Microbiology. 2009 Oct;11(10):2710–9.

59. Gérard E, De Goeyse S, Hugoni M, Agogué H, Richard L, Milesi V, et al. Key Role of Alphaproteobacteria and Cyanobacteria in the Formation of Stromatolites of Lake Dziani Dzaha (Mayotte, Western Indian Ocean). Front Microbiol. 2018 May 22;9:796.

60. Zhu T, Dittrich M. Carbonate Precipitation through Microbial Activities in Natural Environment, and Their Potential in Biotechnology: A Review. Front Bioeng Biotechnol [Internet]. 2016 Jan 20 [cited 2023 Apr 21];4. Available from: http://journal.frontiersin.org/Article/10.3389/fbioe.2016.00004/abstract

61. Brady AL, Slater GF, Omelon CR, Southam G, Druschel G, Andersen DT, et al. Photosynthetic isotope biosignatures in laminated micro-stromatolitic and non-laminated nodules associated with modern, freshwater microbialites in Pavilion Lake, B.C. Chemical Geology. 2010 Jun;274(1–2):56–67.

62. Guo W, Daëron M, Niles P, Goddard WA, Eiler JM. ISOTOPIC FRACTIONATIONS ASSOCIATED WITH DEGASSING OF CO2 FROM AQUEOUS SOLUTIONS AND IMPLICATIONS FOR CARBONATE CLUMPED ISOTOPE THERMOMETRY.

63. Clark ID, Lauriol B. Kinetic enrichment of stable isotopes in cryogenic calcites. Chemical Geology. 1992 Dec;102(1–4):217–28.

64. West JB, Bowen GJ, Dawson TE, Tu KP, editors. Isoscapes [Internet]. Dordrecht: Springer Netherlands; 2010 [cited 2023 Apr 21]. Available from: http://link.springer.com/10.1007/978-90-481-3354-3

65. Han WS, Watson ZT, Kampman N, Grundl T, Graham JP, Keating EH. Periodic changes in effluent chemistry at cold-water geyser: Crystal geyser in Utah. Journal of Hydrology. 2017 Jul;550:54–64.

66. McConnaughey T. 13C and 18O isotopic disequilibrium in biological carbonates: I. Patterns. Geochimica et Cosmochimica Acta. 1989 Jan 1;53(1):151–62.

67. Granger J, Sigman DM, Lehmann MF, Tortell PD. Nitrogen and oxygen isotope fractionation during dissimilatory nitrate reduction by denitrifying bacteria. Limnology & Oceanography. 2008 Nov;53(6):2533–45.

68. Hsieh YJ, Wanner BL. Global regulation by the seven-component Pi signaling system. Current Opinion in Microbiology. 2010 Apr;13(2):198–203.

69. Wanner BL. Phosphorus assimilation and control of the phosphate regulon. In 1996.

70. Lee SY. Bacterial polyhydroxyalkanoates. Biotechnol Bioeng. 1995;49(1):1–14.

71. Patzlaff JS, Van Der Heide T, Poolman B. The ATP/Substrate Stoichiometry of the ATP-binding Cassette (ABC) Transporter OpuA. Journal of Biological Chemistry. 2003 Aug;278(32):29546–51.

72. Shuman HA. Active transport of maltose in Escherichia coli K12. Role of the periplasmic maltose-binding protein and evidence for a substrate recognition site in the cytoplasmic membrane. Journal of Biological Chemistry. 1982 May;257(10):5455–61.

73. Davidson AL, Dassa E, Orelle C, Chen J. Structure, Function, and Evolution of Bacterial ATP-Binding Cassette Systems. Microbiol Mol Biol Rev. 2008 Jun;72(2):317–64.

74. Dippel R, Boos W. The Maltodextrin System of *Escherichia coli* : Metabolism and Transport. J Bacteriol. 2005 Dec 15;187(24):8322–31.

75. Akyel A, Coburn M, Phillips AJ, Gerlach R. Key Applications of Biomineralization. In: Berenjian A, Seifan M, editors. Mineral Formation by Microorganisms [Internet]. Cham: Springer International Publishing; 2022 [cited 2023 Apr 1]. p. 347–87. (Microbiology Monographs; vol. 36). Available from: https://link.springer.com/10.1007/978-3-030-80807-5_10

