## Supplementary Methods for "Meta-omics reveals role of photosynthesis in Microbially Induced Carbonate Precipitation at a CO_2_-rich Geyser"

**Isotope-ratio mass spectrometry**

Organic Carbon

Samples for bulk δ^13^C_org_ analyses were prepared by removing any surficial modern organic material by ultrasonication in 99% methanol. All samples were then reacted with a dilute solution of HCl (2%) at 40°C to remove all carbonate material, and rinsed with deionized water until cleaned. We then dried samples at 80°C and homogenized them before loading them into tin capsules for isotopic analysis using a Costech elemental analyzer attached to a Thermo Delta V isotope ratio mass spectrometer in the Paleo^3^ Laboratory at North Carolina State University. Organic carbon isotope measurements are reported in per mille (‰) relative to the Vienna Pee Dee Belemnite. We used IAEA sucrose (-10.45‰) and caffeine (-27.77‰) standards for calibration. Analytical uncertainty is maintained at < 0.1‰, and replicate analyses had a mean standard error of 0.25‰.

Carbonates

Carbonates were subsampled from collected billets, and cleaned via deionized water before being crushed by hand in a mortar and pestle. Powdered samples were then further cleaned with a dilute H_2_O_2_ (2%) solution to remove organic carbon contamination, rinsed, dried, and analyzed for stable (δ^18^O, δ^13^C) isotopes. Samples were processed in a Nu Carb Automated Carbonate on-line preparation system at 70°C. Powdered samples (400-600 µg) were digested with phosphoric acid (specific gravity 1.94-1.96), and resultant CO_2_ was cryogenically separated after which the evolved CO_2_ was stored in the dual inlet alongside a CO_2_ working gas (δ^13^C= 4.485, δ^18^O= -2.450). Sample δ^18^O and δ^13^C values were measured on a Nu Perspective IS isotope ratio mass spectrometer configured to measure m/z ratios for masses 44–49, and are reported relative to the standard Vienna-Pee Dee Belemnite (VPDB). Solid standards were run before and after each set of eight unknown carbonate samples, and output was referenced to IAEA standards C-1 (δ^13^C= 2.42, δ^18^O= 8.28) and NBS-18 (δ^13^C= -5.014, δ^18^O= -23.2), as well as internal standard C-64 (δ^13^C= -2.05, δ^18^O= -15.54). Data were processed using Easotope software (1) and corrected with a 70°C calcite acid fractionation factor (2). The Pierce Outlier test was used to remove statistical outliers from output (3,4) and the standard error was measured based on a 95% confidence interval; standard error was maintained at <0.25‰ for both carbon and oxygen.

**Scanning electron microscopy and energy-dispersive X-ray spectrometry**

Scanning electron microscopy was done at the Analytical Instrumentation Facility at North Carolina State University. We used the same powdered carbonates that were prepared for isotope ratio mass spectrometry (see above). Carbonates were imaged with a Hitachi SU3900 (Shaumburg, IL, USA) scanning electron microscope. Elemental analysis of a selected area was performed with an energy-dispersive X-ray spectrometer (Oxford Ultim-max, Concord, MA, USA) with 20 keV beam energy. The Ultimax is a silicon drift detector with a 40 mm^2^ area. Elements were quantified using Oxford AZtec software. The sum of net amounts of detected elements were normalized to 100% wt%; average percentages of the individual compounds and standard deviations from the mean were calculated across all images. Only elements with 0.5% abundance or higher were considered for our analyses.

**Assembly, binning and annotation of metagenomes**

Prokaryotic sequences

We extracted all prokaryotic contigs identified with Whokaryote from the metagenomic assemblies using the filterbyname.sh implemented in bbmap (29). We mapped all raw reads to these contigs with bbmap to use mapping coverages for binning. Contigs were binned using MetaBat2 (5). We obtained 197 bins for CG-1 travetine and 343 bins for CG-10 stravertine. We checked bin quality with checkM (6) and checkM2 (7), and bin rRNA and tRNA content with barrnap (8) and aragorn (9). We corrected contamination scores reported with checkM2 with the strain heterogeneity values obtained from checkM to exclude strain level contamination from the contamination values. We used the prodigal gene predictions obtained from checkM2 and functionally annotated them with Microbeannotator (10). We manually refined protein annotations for our proteins of interest using InterProScan (11) and Uniprot Blast against the Swissprot database (12).

Eukaryotic sequences

We extracted all eukaryotic contigs identified with Whokaryote as described above. We then predicted open reading frames (ORFs) with BUSCO (13) using the following parameters: -m geno --augustus --auto-lineage-euk. We concatenated all predicted genes files and considered them as one eukaryotic “bin”.

**Protein extraction, peptide preparation, and determination**

We extracted proteins in triplicate per sampling site. We prepared tryptic peptides by combining a trichloroacetic acid (TCA) precipitation protocol adapted from Qian and Hettich (14) and the filter-aided-sample-preparation (FASP) protocol (15) (Suplementary Methods). We suspended 5 g of travertine homogenate in 15 mL SDT lysis buffer (4% [wt/vol] SDS, 100 mM Tris-HCl, pH 7.6, 0.1 M DTT) and sonicated with a microtip probe on a QSonica Q700 sonicator using 4 cycles of 25 s at 20% amplitude (700 Watt) with 25 s dwell time between cycles, followed by heating at 95°C for 10 min. After heating, we centrifuged the samples for 10 min at 15 000 x g and transferred the supernatant into a fresh 50 mL conical tube following another centrifugation cycle for 5 min at 15 000 x g. We again transferred the supernatant into a fresh conical tube and added 15 mL of 20% TCA solution prepared in ice cold acetone resulting in a final TCA concentration of 10% (w/v). The mixture was briefly vortexed and stored at -80°C for overnight precipitation. The next day, we thawed the samples and pelleted proteins by centrifugation at 15,000 × g at 4°C for 20 min. We discarded the supernatant and washed the pellet once with 100% cold acetone and twice with 80% cold acetone following another precipitation in 80% acetone at -20°C for 20 minutes. Following the precipitation, we centrifuged the samples for 15,000 × g at 4°C for 15 min. We discarded the supernatant and washed the pellet with 80% cold acetone. We air dried the pellet and resuspended it in SDT buffer, followed by heating at 95°C for 10 min. Subsequently, we followed the FASP protocol with the modification that we did not add additional UA to the suspended protein pellet to avoid further sample dilution when loading on the 10-kDa-molecular-weight-cutoff (MWCO) 500-μl centrifugal filter units (VWR International). We centrifuged the lysate at 14,000 × g for 40 min and added 200 μl of UA solution and centrifuged the mixture again at 14,000 × g for 40 min. We added 100 μl of IAA solution (0.05 M iodoacetamide in UA solution) and then incubated the samples at 22°C for 20 min in the dark. We removed the IAA solution by centrifugation, followed by three wash steps with 100 μl of UA solution. Subsequently, we washed the filters three times with 100 μl of ABC buffer (50 mM ammonium bicarbonate). We added 0.55 μg of Pierce mass spectrometry (MS)-grade trypsin (Thermo Fisher Scientific) in 40 μl of ABC buffer to each filter. Filters were incubated overnight in a wet chamber at 37°C. The next day, we eluted the peptides by centrifugation at 14,000 × g for 20 min, followed by a second elution with of 50 μl of 0.5 M NaCl. Peptides were quantified using the Pierce micro-bicinchoninic acid (microBCA) kit (Thermo Fisher Scientific), following the instructions of the manufacturer.

**Protein identification and quantification**

For protein identification, we built a custom protein sequence database for each sampling site using all ORFs predicted from eukaryotic and prokaryotic bins as well as the unassigned contigs that were filtered out by Whokaryote. All predicted sequences were concatenated into one database per sampling site. We then clustered each database with CD-HIT (16) using 95% sequence identity (17). We appended the cRAP protein sequence database (http://www.thegpm.org/crap) of common laboratory contaminants to each database. The database contained 1,595,960 protein sequences for CG-1 and 3,887,042 protein sequences for CG-10. We performed searches of the MS/MS spectra against this database with the Sequest HT node in Proteome Discoverer version 2.3.0.388 (Thermo Fisher Scientific), as described by Jensen et al. (18). The following parameters were used: trypsin (full), maximum of 2 missed cleavages, 10 ppm precursor mass tolerance, 0.1 Da fragment mass tolerance, and maximum of 3 equal dynamic modifications per peptide, namely, oxidation on M (+15.995 Da), carbamidomethyl on C (+57.021 Da), and acetyl on the protein N terminus (+42.011 Da). FDRs for peptide spectral matches (PSMs) were calculated and filtered using the Percolator node in Proteome Discoverer (52). Percolator was run with a maximum Delta CN of 0.05, a strict target FDR of 0.01, a relaxed target FDR of 0.05, and validation based on q-value. The Protein FDR Validator node in Proteome Discoverer was used to calculate q-values for inferred proteins based on the results from a search against a target-decoy database. Proteins with q-values of <0.01 were categorized as high-confidence identifications, and proteins with q-values of 0.01 to 0.05 were categorized as medium-confidence identifications. We combined search results for all samples into a multiconsensus report in Proteome Discoverer, and only Master proteins identified with medium or high confidence were retained, resulting in an overall protein-level FDR of 5%. For protein quantification, normalized spectral abundance factors (NSAFs) (19) were calculated and multiplied by 100, to give the relative protein abundance in %.

**References**

1.	John CM, Bowen D. Community software for challenging isotope analysis: First applications of ‘Easotope’ to clumped isotopes: Community software for challenging isotope analysis. Rapid Commun Mass Spectrom. 2016 Nov 15;30(21):2285–300.

2.	Kim ST, O’Neil JR. Equilibrium and nonequilibrium oxygen isotope effects in synthetic carbonates. Geochimica et Cosmochimica Acta. 1997 Aug;61(16):3461–75.

3.	Huntington KW, Eiler JM, Affek HP, Guo W, Bonifacie M, Yeung LY, et al. Methods and limitations of ‘clumped’ CO _2_ isotope (Δ _47_ ) analysis by gas-source isotope ratio mass spectrometry. J Mass Spectrom. 2009 Sep;44(9):1318–29.

4.	Burgener L, Huntington KW, Hoke GD, Schauer A, Ringham MC, Latorre C, et al. Variations in soil carbonate formation and seasonal bias over >4 km of relief in the western Andes (30°S) revealed by clumped isotope thermometry. Earth and Planetary Science Letters. 2016 May;441:188–99.

5.	Kang DD, Li F, Kirton E, Thomas A, Egan R, An H, et al. MetaBAT 2: an adaptive binning algorithm for robust and efficient genome reconstruction from metagenome assemblies. PeerJ. 2019 Jul 26;7:e7359.

6.	Parks DH, Imelfort M, Skennerton CT, Hugenholtz P, Tyson GW. CheckM: assessing the quality of microbial genomes recovered from isolates, single cells, and metagenomes. Genome Res. 2015 Jul;25(7):1043–55.

7.	Chklovski A, Parks DH, Woodcroft BJ, Tyson GW. CheckM2: a rapid, scalable and accurate tool for assessing microbial genome quality using machine learning [Internet]. Bioinformatics; 2022 Jul [cited 2023 Apr 21]. Available from: http://biorxiv.org/lookup/doi/10.1101/2022.07.11.499243

8.	Seemann, Torsten. Barrnap 0.7: rapid ribosomal RNA prediction [Internet]. 2013. Available from: https://github.com/tseemann/barrnap

9.	Laslett D. ARAGORN, a program to detect tRNA genes and tmRNA genes in nucleotide sequences. Nucleic Acids Research. 2004 Jan 2;32(1):11–6.

10.	Ruiz-Perez CA, Conrad RE, Konstantinidis KT. MicrobeAnnotator: a user-friendly, comprehensive functional annotation pipeline for microbial genomes. BMC Bioinformatics. 2021 Dec;22(1):11.

11.	Jones P, Binns D, Chang HY, Fraser M, Li W, McAnulla C, et al. InterProScan 5: genome-scale protein function classification. Bioinformatics. 2014 May 1;30(9):1236–40.

12.	The UniProt Consortium, Bateman A, Martin MJ, Orchard S, Magrane M, Ahmad S, et al. UniProt: the Universal Protein Knowledgebase in 2023. Nucleic Acids Research. 2023 Jan 6;51(D1):D523–31.

13.	Simão FA, Waterhouse RM, Ioannidis P, Kriventseva EV, Zdobnov EM. BUSCO: assessing genome assembly and annotation completeness with single-copy orthologs. Bioinformatics. 2015 Oct 1;31(19):3210–2.

14.	Qian C, Hettich RL. Optimized Extraction Method To Remove Humic Acid Interferences from Soil Samples Prior to Microbial Proteome Measurements. J Proteome Res. 2017 Jul 7;16(7):2537–46.

15.	Wiśniewski JR, Zougman A, Nagaraj N, Mann M. Universal sample preparation method for proteome analysis. Nat Methods. 2009 May;6(5):359–62.

16.	Fu L, Niu B, Zhu Z, Wu S, Li W. CD-HIT: accelerated for clustering the next-generation sequencing data. Bioinformatics. 2012 Dec;28(23):3150–2.

17.	Blakeley-Ruiz JA, Kleiner M. Considerations for constructing a protein sequence database for metaproteomics. Computational and Structural Biotechnology Journal. 2022;20:937–52.

18.	Jensen M, Wippler J, Kleiner M. Evaluation of RNA *later* as a Field-Compatible Preservation Method for Metaproteomic Analyses of Bacterium-Animal Symbioses. Gralnick JA, editor. Microbiol Spectr. 2021 Oct 31;9(2):e01429-21.

19.	Zybailov B, Mosley AL, Sardiu ME, Coleman MK, Florens L, Washburn MP. Statistical Analysis of Membrane Proteome Expression Changes in *Saccharomyces* *c* *erevisiae*. J Proteome Res. 2006 Sep 1;5(9):2339–47.
