## Supplementary Material for "Meta-omics reveals role of photosynthesis in Microbially Induced Carbonate Precipitation at a CO_2_-rich Geyser"

**Supplementary figures and tables**

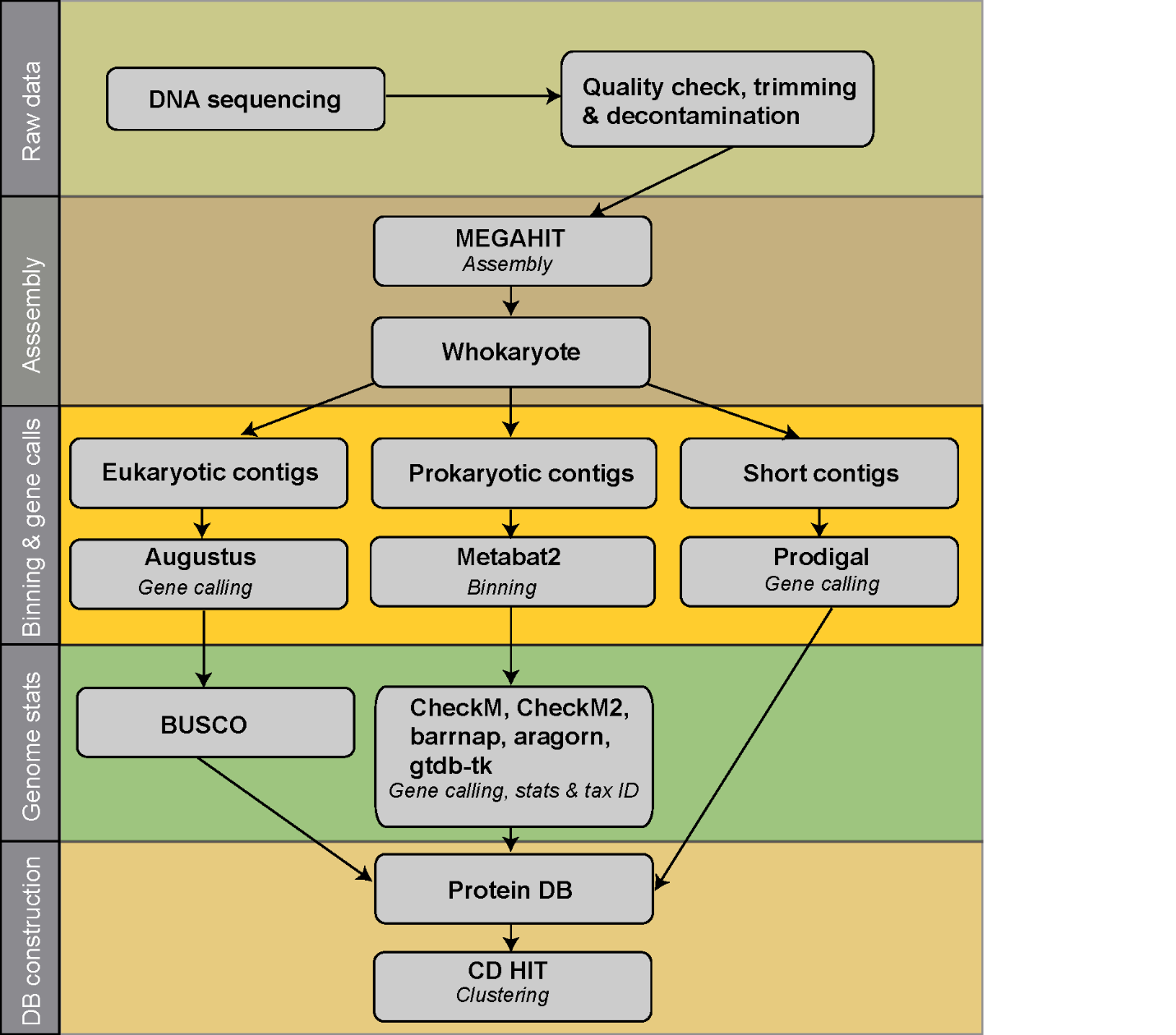

**Figure S1:** Overview of the metagenomics workflow used for data processing and protein database curation.

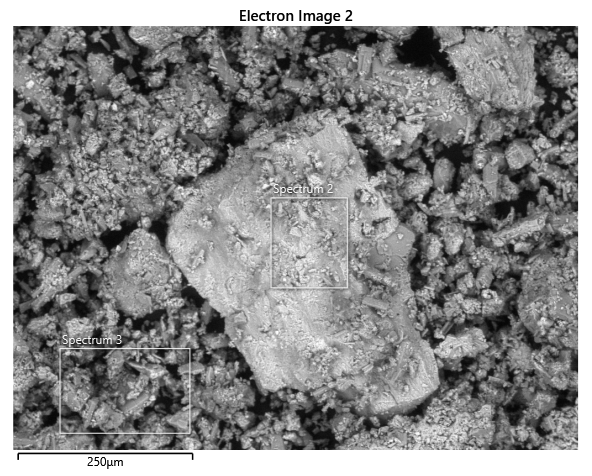

**Figure S2:** Scanning electron microscopy (SEM) photomicrographs and energy dispersive X-ray (EDX) analyses of the carbonate minerals present in travertine collected from CG-1**.** The analysis was performed with a Hitachi SU3900 scanning electron microscope. Elemental analysis of a selected area was performed with an energy-dispersive X-ray spectrometer with 20 keV beam energy. Elements were quantified using Oxford AZtec software. Elemental quantification of the measured spectra can be found in Table S1.

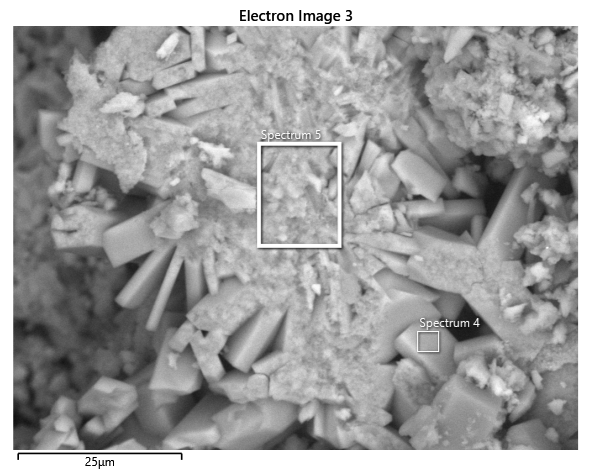

**Figure S3:** SEM photomicrographs and EDX analyses of the carbonate minerals present in travertine collected from CG-1**.** The analysis was performed with a Hitachi SU3900 scanning electron microscope. Elemental analysis of a selected area was performed with an energy-dispersive X-ray spectrometer with 20 keV beam energy. Elements were quantified using Oxford AZtec software. Elemental quantification of the measured spectra can be found in Table S1.

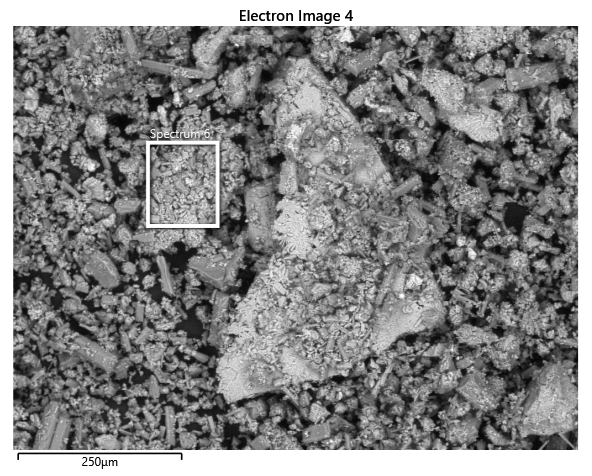

**Figure S4:** SEM photomicrographs and EDX analyses of the carbonate minerals present in travertine collected from CG-1**.** The analysis was performed with a Hitachi SU3900 scanning electron microscope. Elemental analysis of a selected area was performed with an energy-dispersive X-ray spectrometer with 20 keV beam energy. Elements were quantified using Oxford AZtec software. Elemental quantification of the measured spectra can be found in Table S1.

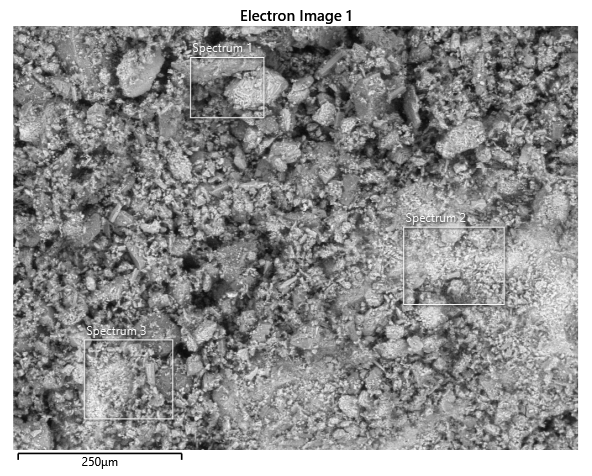

**Figure S5:** SEM photomicrographs and EDX analyses of the carbonate minerals present in travertine collected from CG-10**.** The analysis was performed with a Hitachi SU3900 scanning electron microscope. Elemental analysis of a selected area was performed with an energy-dispersive X-ray spectrometer w ith 20 keV beam energy. Elements were quantified using Oxford AZtec software. Elemental quantification of the measured spectra can be found in Table S1.

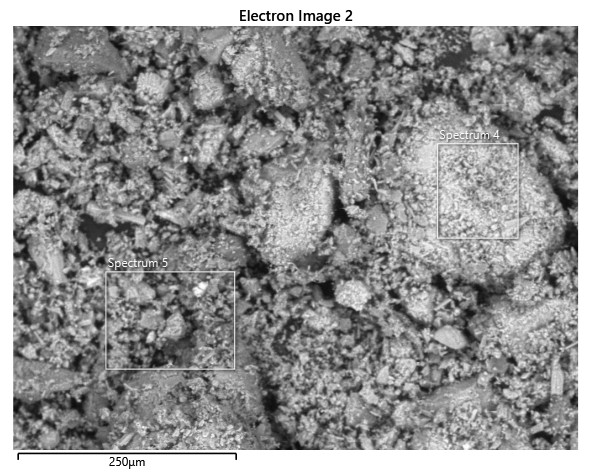

**Figure S6:** SEM photomicrographs and EDX analyses of the carbonate minerals present in travertine collected from CG-10. The analysis was performed with a Hitachi SU3900 scanning electron microscope. Elemental analysis of a selected area was performed with an energy-dispersive X-ray spectrometer with 20 keV beam energy. Elements were quantified using Oxford AZtec software. Elemental quantification of the measured spectra can be found in Table S1.

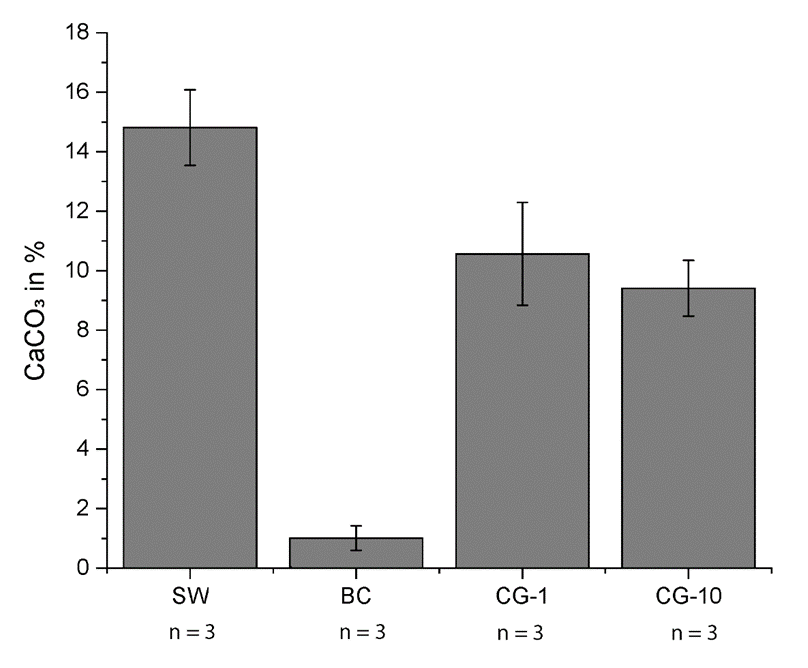

**Figure S7:** Carbonate quantification of CG travertine collected at CG-1 and CG-10. SW are stromatolite samples collected from Rife Bed, Wyoming and BC are soil samples from Cedar Mountain Base Camp, Utah. These samples were included as references as they are known to contain large and small amounts of CaCO_3_ respectively. Error bars indicate the standard deviation of the mean across 3 replicates. Quantities are displayed in g of CaCO_3_ per input sediment weight in % (wt/wt).

**Table S1:** EDX analyses of the carbonate minerals present in travertine collected from CG-1. Only elements that are above the detection limit threshold of 0.5% are listed. Spectrum 1 was discarded due to calibration of the instrument. The analysis was performed with a Hitachi SU3900 scanning electron microscope. Elemental analysis of a selected area was performed with an energy-dispersive X-ray spectrometer with 20 keV beam energy. Elements were quantified using Oxford AZtec software.

| Element | Spectrum1 | Spectrum2 | Spectrum3 | Spectrum4 | Spectrum5 | Spectrum6 |
| --- | --- | --- | --- | --- | --- | --- |
| O | Invalid | 43.4 | 43.8 | 45.7 | 44.7 | 44.4 |
| Ca | Invalid | 27 | 23 | 34.8 | 35.6 | 23.3 |
| C | Invalid | 14 | 15 | 14.4 | 13.5 | 14.5 |
| S | Invalid | 5.6 | 7.2 | 2.8 | 2.5 | 7.8 |
| N | Invalid | 4.3 | 4.9 | 0 | 1.8 | 5.4 |
| Fe | Invalid | 3.4 | 4.5 | 1.5 | 1.5 | 3.5 |
| Si | Invalid | 1.4 | 0.6 | 0 | 0 | 0 |

**Table S2:** EDX analyses of the carbonate minerals present in travertine collected from CG-10. Only elements that are above the detection limit threshold of 0.5% are listed. The analysis was performed with a Hitachi SU3900 scanning electron microscope. Elemental analysis of a selected area was performed with an energy-dispersive X-ray spectrometer with 20 keV beam energy. Elements were quantified using Oxford AZtec software.

| Element | Spectrum1 | Spectrum2 | Spectrum3 | Spectrum4 | Spectrum5 |
| --- | --- | --- | --- | --- | --- |
| O | 41.8 | 45.2 | 44.2 | 43.4 | 46.2 |
| Ca | 33.4 | 31.1 | 29.2 | 25.9 | 26.9 |
| C | 13.6 | 14.2 | 12.9 | 15.5 | 15.9 |
| S | 4.1 | 4.6 | 6 | 9.3 | 6.6 |
| N | 3.6 | 1.3 | 3.4 | 2.6 | 0 |
| Fe | 1.8 | 1.3 | 1.9 | 1 | 2 |
| Mg | 0.5 | 0 | 0.8 | 0 | 0.6 |
| Na | 0.5 | 0.9 | 0.6 | 1.3 | 0.7 |

**Table S3:** IRMS of organic carbon and nitrogen derived from travertine samples collected at CG. Samples were measured with a Costech elemental analyzer attached to a Thermo Delta V isotope ratio mass spectrometer. Organic carbon isotope measurements are reported in per mille (‰) relative to the Vienna Pee Dee Belemnite (VPDB).

| Sample ID | Weight (mg) | mg C | % C per sample | mg N | % N per sample | δ^13^C_VPDB_ | δ^15^N_VPDB_ | C:N |
| --- | --- | --- | --- | --- | --- | --- | --- | --- |
| CG-1 | 85 | 0.350 | 0.41 | 0.070 | 0.08 | -25.5 | 3.0 | 5.8 |
| CG-10 | 61 | 0.152 | 0.25 | 0.031 | 0.05 | -21.5 | 3.7 | 5.7 |

**Table S4:** Mass of CaCO3 (wt%) results of gasometric CaCO_3_ quantification of travertine samples collected at CG-1 and CG-10. Stromatolite carbonates (SW) were measured as a control for high CaCO_3_ content and sandy soils (BC) were measured as a control for low CaCO_3_ content.

| Samples | Weight [g] | CaCO3 [g] | %CaCO3 |
| --- | --- | --- | --- |
| SW-1 | 13.02 | 2.05 | 15.73 |
| SW-2 | 13 | 1.74 | 13.36 |
| SW-3 | 13.34 | 2.05 | 15.35 |
| BC-1-1 | 13.21 | 0.14 | 1.02 |
| BC-1-2 | 13.4 | 0.19 | 1.41 |
| BC-1-3 | 12.99 | 0.08 | 0.59 |
| CG-1-1 | 13.78 | 1.37 | 9.96 |
| CG-1-2 | 13.66 | 1.26 | 9.22 |
| CG-1-3 | 13.3 | 1.67 | 12.52 |
| CG-10-1 | 13.42 | 1.13 | 8.38 |
| CG-10-2 | 12.11 | 1.24 | 10.22 |
| CG-10-3 | 13.1 | 1.26 | 9.62 |

**Table S5**: Determination of significant differences in terms of CaCO_3_ amount between CG travertine samples and the controls. Statistical testing was performed using a two-tailed *Student’s t-test*. P-values below 0.05 are considered significant.

| Comparison | p-value |
| --- | --- |
| CG-1:CG-10 | 0.36 |
| CG-1:SW | 0.03 |
| CG-1:BC | 0.00 |
| CG-10:SW | 0.00 |
| CG-10:BC | 0.00 |

**Table S6** – Comparison of mappable metagenomics reads between CG-1 and CG-10. Reads were mapped to co-assemblies using libraries of both sequencing facilities. Mapping was done with bbmap.

| Mapping | Total Reads used | Read 1 Mapped | Read 2 Mapped | Mapped reads | Mapped % | Unmapped | Unmapped% |
| --- | --- | --- | --- | --- | --- | --- | --- |
| CG-1 (GSL) | 228767908 | 111823910 | 111357839 | 223181749 | 97.56 | 5586159 | 2.44 |
| CG-1 (BGI) | 221518738 | 108010629 | 107996773 | 216007402 | 97.51 | 5511336 | 2.49 |
| CG-10 (GSL) | 218793910 | 102553682 | 102069386 | 204623068 | 93.52 | 14170842 | 6.48 |
| CG-10 (BGI) | 220631020 | 102144086 | 102102308 | 204246394 | 92.57 | 16384626 | 7.43 |

**Table S7 -** Isotope ratio mass spectrometry measurements (IRMS) of carbonates derived from travertine samples collected at CG**.** Sample δ^18^O and δ^13^C values were measured on a Nu Perspective IS isotope ratio mass spectrometer configured to measure m/z ratios for masses 44–49. Isotope measurements are reported in per mille (‰) relative to the Vienna Pee Dee Belemnite (VPDB) standard and the Vienna Standard Mean Ocean Water (VSMOW) respectively.

| Replicate *#* | Analysis | Site | δ^13^C _VPDB_ | δ^18^O _VPDB_ | δ^18^O _VSMOW_ |
| --- | --- | --- | --- | --- | --- |
| R1 | CO_2_ bulk | CG-1 | 7.49 | -12.21 | 18.33 |
| R2 | CO_2_ bulk | CG-1 | 7.27 | -12.46 | 18.07 |
| R3 | CO_2_ bulk | CG-1 | 7.19 | -12.34 | 18.2 |
| R1 | CO_2_ bulk | CG-10 | 7.41 | -12.36 | 18.18 |
| R2 | CO_2_ bulk | CG-10 | 7.37 | -12.38 | 18.15 |
| R3 | CO_2_ bulk | CG-10 | 7.36 | -12.46 | 18.07 |
